# Three-Component Model of the Spinal Nerve Branching Pattern, based on the View of the Lateral Somitic Frontier and Experimental Validation

**DOI:** 10.1101/2020.07.29.227710

**Authors:** Shunsaku Homma, Takako Shimada, Ikuo Wada, Katsuji Kumaki, Noboru Sato, Hiroyuki Yaginuma

## Abstract

One of the decisive questions about human gross anatomy is unmatching the adult branching pattern of the spinal nerve to the embryonic lineages of the peripheral target muscles. The two principal branches in the adult anatomy, the dorsal and ventral rami of the spinal nerve, innervate the intrinsic back muscles (epaxial muscles), as well as the body wall and appendicular muscles (hypaxial muscles), respectively. However, progenitors from the dorsomedial myotome develop into the back and proximal body wall muscles (primaxial muscles) within the sclerotome-derived connective tissue environment. In contrast, those from the ventrolateral myotome develop into the distal body wall and appendicular muscles (abaxial muscles) within the lateral plate-derived connective tissue environment. Thus, the ventral rami innervate muscles that belong to two different embryonic compartments. Because strict correspondence between an embryonic compartment and its cognate innervation is a way to secure the development of functional neuronal circuits, this mismatch indicates that we may need to reconcile our current understanding of the branching pattern of the spinal nerve with regard to embryonic compartments. Accordingly, we first built a model for the branching pattern of the spinal nerve, based on the primaxial-abaxial distinction, and then validated it using mouse embryos.

In our model, we hypothesized the following: 1) a single spinal nerve consists of three nerve components: primaxial compartment-responsible branches, a homologous branch to the canonical intercostal nerve bound for innervation to the abaxial compartment in the ventral body wall, and a novel class of nerves that travel along the lateral cutaneous branch to the appendicles; 2) the three nerve components are discrete only during early embryonic periods but are later modified into the elaborate adult morphology; and 3) each of the three components has its own unique morphology regarding trajectory and innervation targets. Notably, the primaxial compartment-responsible branches from the ventral rami have the same features as the dorsal rami. Under the above assumptions, our model comprehensively describes the logic for innervation patterns when facing the intricate anatomy of the spinal nerve in the human body.

In transparent whole-mount specimens of embryonic mouse thoraces, the single thoracic spinal nerve in early developmental periods trifurcated into superficial, deep, and lateral cutaneous branches; however, it later resembled the adult branching pattern by contracting the superficial branch. The superficial branches remained segmental while the other two branches were free from axial restriction. Injection of a tracer into the superficial branches of the intercostal nerve labeled Lhx3-positive motoneurons in the medial portion of the medial motor column (MMCm). However, the injection into the deep branches resulted in retrograde labeling of motoneurons that expressed Oct6 in the lateral portion of the medial motor column (MMCl). Collectively, these observations on the embryonic intercostal nerve support our model that the spinal nerve consists of three distinctive components.

We believe that our model provides a framework to conceptualize the innervation pattern of the spinal nerve based on the distinction of embryonic mesoderm compartments. Because such information about the spinal nerves is essential, we further anticipate that our model will provide new insights into a broad range of research fields, from basic to clinical sciences.

## INTRODUCTION

The peripheral branching pattern of the spinal nerve constitutes an essential chapter of the human gross anatomy, and has long been a subject of study in basic and clinical medicine. Nevertheless, a fundamental puzzle has remained: the branching pattern in the adult anatomy does not reflect the embryonic lineages of the peripheral target muscles.

In the adult, once it leaves the vertebrae, the spinal nerve bifurcates into the dorsal and ventral rami. The dorsal ramus innervates the back muscles, whereas the ventral ramus innervates the body wall and appendicular muscles (Standring, 2015). The terms of ‘epaxial’ and ‘hypaxial’ have been used to describe each respective target of motor innervations by the dorsal and ventral rami (Spörle, 2001). However, the ‘epaxial’ and ‘hypaxial’ distinction does not correspond to embryonic somitic lineages (Burke and Nowicki, 2003). The medial portion of the somite gives rise to the back muscles and some body wall muscles, such as the proximal intercostal muscles, whereas the lateral portion gives rise to the distal body wall muscles and appendicular muscles (Ordahl and Le Douarin, 1992; Olivera-Martinez et al., 2000). Two muscle lineages develop exclusively in different connective tissue environments, derived either from the sclerotome or from the lateral plate mesoderm (LPM). The term ‘primaxial’ is used to describe the muscles that are derived from the medial somite and are differentiated in the sclerotome-derived connective tissue environment, whereas ‘abaxial’ for muscles from the lateral somite and in the LPM-derived connective tissue environment (Burke and Nowicki et al., 2003). The boundary between the primaxial and abaxial muscle-containing compartments is named the lateral somitic frontier (Nowicki et al., 2003; Burke and Nowicki, 2003; Durland et al., 2008). Thus, the ventral ramus innervates muscles located before and beyond the lateral somitic frontier, whereas the dorsal ramus of the spinal nerve innervates muscles only before the frontier. Hence, the dorsal and ventral rami of the spinal nerve in adult anatomy is thought of as a secondary modification on an initial differential innervation laid down during early developmental periods (Gilbert and Barresi, 2016). However, an original innervation pattern based on the primaxial-abaxial distinction has not yet been investigated, and its development into the adult morphology has yet to be analyzed.

As the muscles have two different lineages, the target connective tissue of sensory innervation, the dermis, also has two separate embryonic origins (Mauger, 1972; Durland et al., 2008; Fliniaux et al., 2004). The dermis in the primaxial and abaxial compartments are from the somite and the LPM, respectively. Dual sources of the dermis inevitably bring us the second related puzzle of how a single sensory dermatome in the body wall receives coherent sensory innervations. The axial specification by the Hox gene-codes is different between the paraxial and lateral plate mesoderm (Cohn et al., 1997; Nowicki and Burk, 2000). However, current dermatome maps employ the axial levels of paraxial somites, from which the dermis is derived, even for the abaxial compartments. The incompatible axial specification of the dermis and spinal nerve brings into puzzle the global alignment of the spinal nerve with the lateral structures.

The third puzzle is how the columnar organization of spinal motoneurons reflects embryonic innervation patterns to muscles in the primaxial and abaxial compartments. The motoneurons in the spinal cord are organized into two primary columns, the medial motor column (MMC) and lateral motor column (LMC). The MMC is further subdivided into medial (MMCm) and lateral (MMCl) portions (Callister et al., 1987; Gutman et al., 1993; Kitamura and Richmond, 1994; Vanderhorst and Holstege, 1997; Watson, 2009). MMCm motoneurons innervate the back muscles through the dorsal rami, whereas the MMCl and LMC motoneurons innervate the body wall and appendicular muscles through the ventral rami. This columnar distinction was initially prescribed based on the locations of a specific motoneuron group to innervate a particular muscle (motoneuron pool) within the ventral horn. Later, the combinatorial expression patterns of the LIM-class homeobox transcription factors were found to coincide with the columnar organization (Tsuchida et al., 1994; Jessel, 2000). For this reason, the current understanding of the columnar organization does not incorporate the concept of the lateral somitic frontier from the start. Furthermore, MMCm motoneurons send axons to some paraxial muscles through the ventral rami (Stillhard, 1981; Luxenhofer et al., 2014), and the motoneurons innervating the shoulder girdle muscle share a molecular signature with MMCm motoneurons (e.g., rhomboid muscle; Tsuchida et al., 1994). These exemptions for the setting on the MMCm further imply that we may need to accommodate the current mapping of MMC motoneurons to targets with an alternative classification scheme of peripheral muscles.

The ventral ramus of the thoracic spinal nerve, namely the intercostal nerve, has been thought of as a representative example of the spinal nerve. The typical intercostal nerves (the second to eleventh) are of single cord in shape, running along the lower margin of the costae and sequentially sending out twigs to nearby muscles such as the intercostal muscles (Standring, 2015). The intricate branching patterns of plexuses in the other spinal levels have been generally perceived as an elaboration from the simple segmental design of the intercostal nerve, as the varieties of target muscles increase in the girdle and limb. However, the logic of this elaboration has never been described: therefore, it is the fourth puzzle in the present study. Because the extent of this elaboration is proportional to the amount of LPM-associated targets, such as the limb, this logic may be a manifestation of the lateral somitic frontier for the branching pattern of the spinal nerve. Conversely, attaining this logic may eventually lead us to a comprehensive classification of all the branches of the spinal nerve, and hence a better understanding of the innervation pattern in the human body.

To answer the above four puzzles, we first constructed a model for the spinal nerve branching pattern, which is compatible with the concept of the lateral somitic frontier, and explored it in human anatomy. We then validated our model in the mouse embryos, by analyzing the gross morphology of the spinal nerve and the expression pattern of molecular markers in motoneurons after retrograde labeling.

## RESULTS and DISCUSSIONS

### 1. Three-component model

Our model theorizes that a single ventral ramus of the spinal nerve is a composite that consists of three elements: 1) branches to innervate body wall muscles and dermis in the primaxial compartment; 2) an outer body wall branch, which travels along with the lateral cutaneous branch, to convey motor and sensory innervations to the appendicle in the abaxial compartment; and 3) an additional branch to enforce the innervation to ventral body wall muscles and dermis in the abaxial compartment (Figures 1 and 2). The three primary branches are independent only in early developmental periods. However, they are later either modified into a seemingly single cord, as in the case of the intercostal nerve, or fabricated into plexuses in other axial levels so that only terminal segments can be recognized as twigs or higher-ordered branches to a motor and sensory target, as observed in the adult. The autonomic nervous system was excluded from our model because the significant target structures are smooth muscle, and were hence not subject to primaxial and abaxial dichotomy.

**Figure 1.**
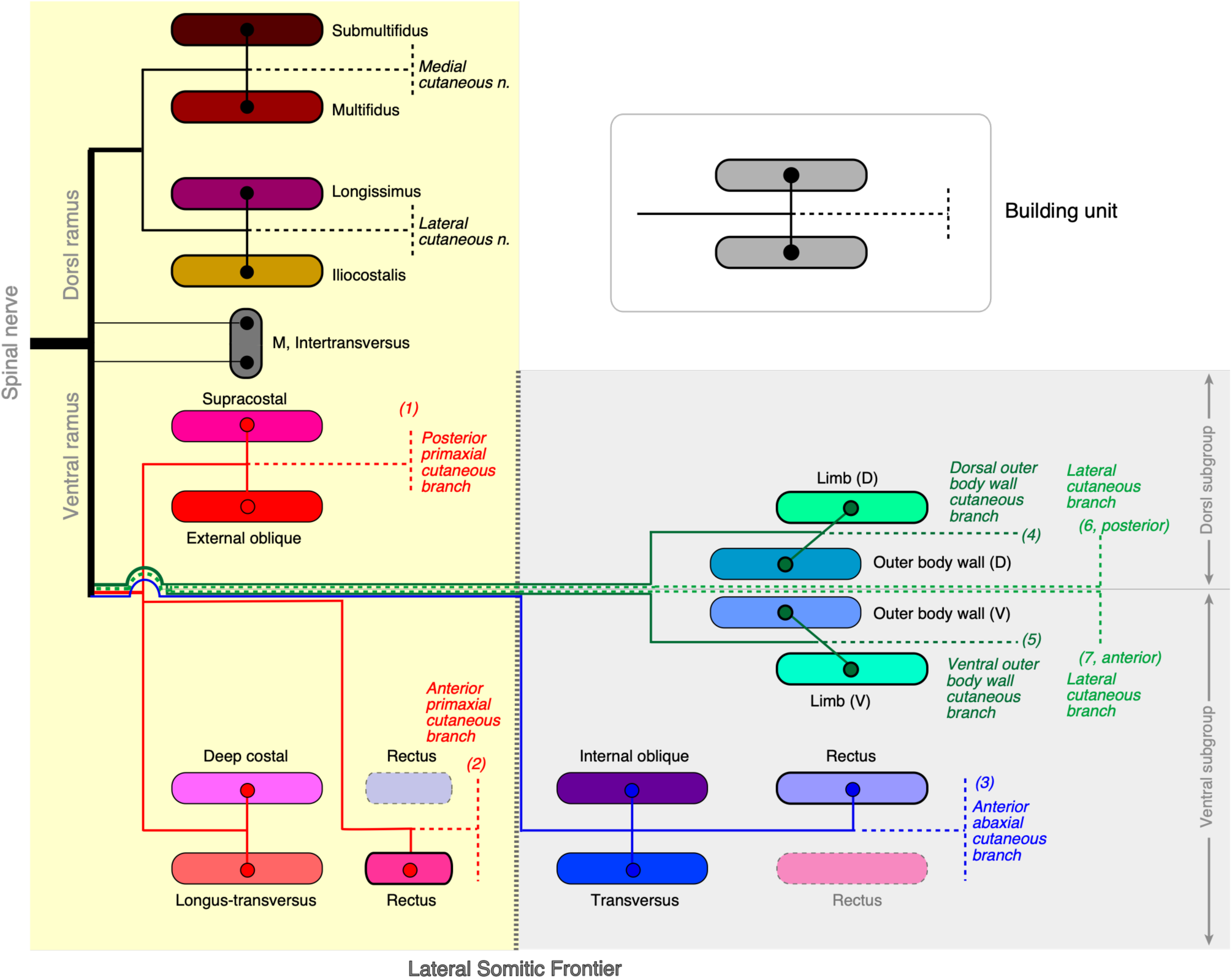
Diagram of mapping the three nerve components to muscle classes. Primaxial compartment-responsible branches are the red lines. Branches homologous to the canonical intercostal nerve (abaxial compartment-responsible) are the blue lines. The lateral cutaneous branch is the light green line, and its accompanied branches to muscles on the body wall and in the appendicles are the dark green lines. The hatched lines are the extension of cutaneous branches from the parental muscular branches. The building unit used in the model construction is in the upper-right square. However, one member of each pair of rectus muscles is absent in the human body and is illustrated in the opaque rectangles. The primaxial compartment is in the cream yellow-colored box, and the abaxial compartment is in the light gray-colored box.

**Figure 2.**
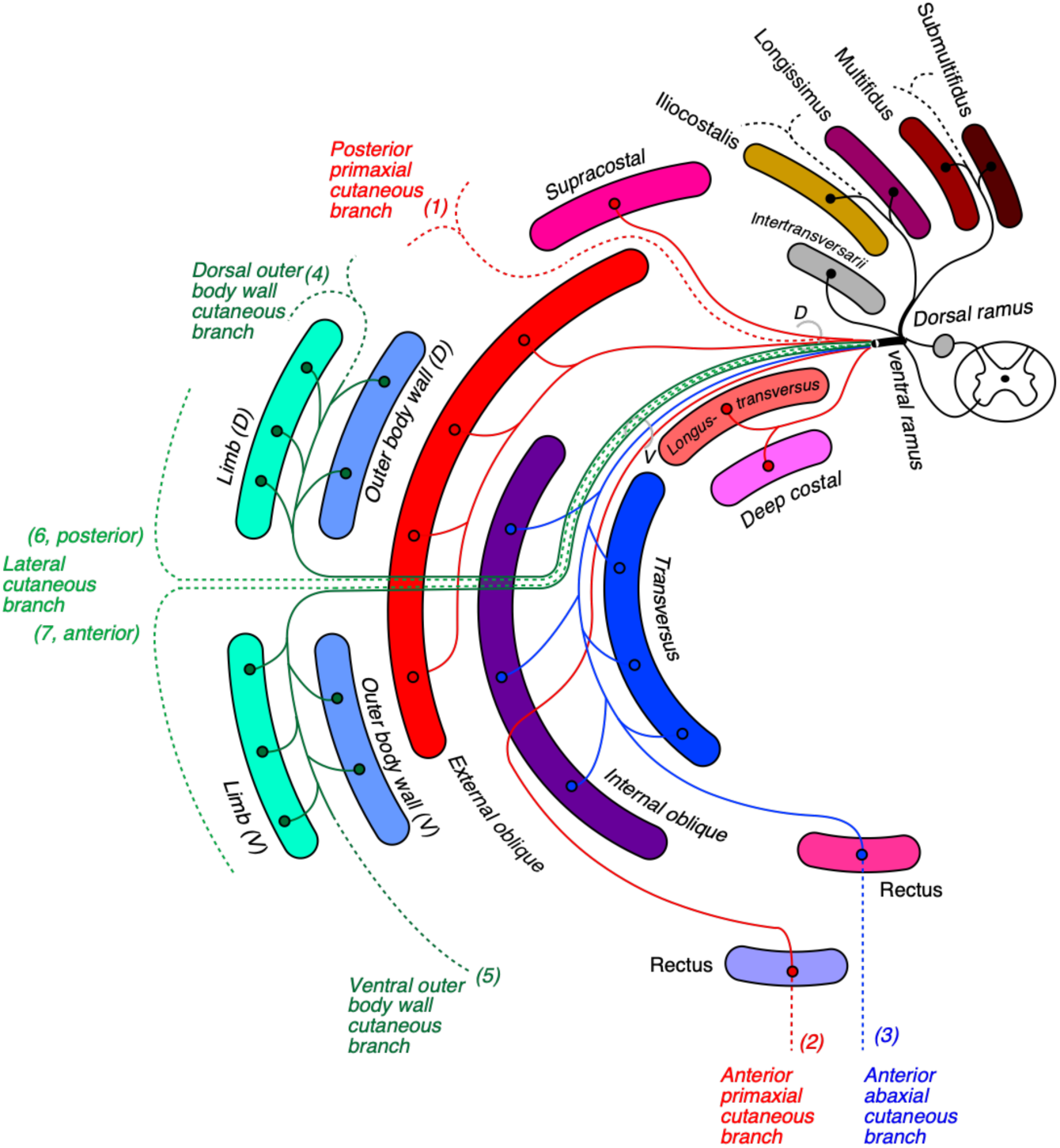
Diagram of nerve-muscle mapping with layer information. Note that muscle layers in the primaxial compartment of the body wall are four in number (deep costal, longus-transvesus, external oblique, and supracostal), and that limb muscles (outer body wall and limb classes) are entirely independent of the layer organization of the body wall. Three nerve components spread out into different intermuscular spaces. The canonical intercostal nerve or its homolog (blue) runs in the deep intermuscular space, and primaxial compartment-responsible nerves (red) run in the superficial intermuscular space. The lateral cutaneous and outer body wall branches (light and dark green) travel outside of the body wall. There are seven presumed seven cutaneous branches: (1) the posterior primaxial cutaneous branch (red hatched line); (2) the anterior cutaneous branch in the primaxial compartment (red hatched line); (3) the anterior abaxial cutaneous branch (blue hatched line); (4 and 5) the dorsal and ventral outer body wall cutaneous branches (hatched, dark green lines); and (6 and 7) the lateral cutaneous branches (hatched, light green lines). Nerve branches that innervate muscle in the dorsal and ventral subgroups are in bundles of (D) and (V), respectively.

When we created the above model, we first made a working hypothesis that the branches from the ventral ramie to innervate the muscles that are within the primaxial compartment also possess the same traits shared by the dorsal ramie, because the factors that determine where branches diverge from are likely to reflect the spatial and temporal factors of nerve-muscle interactions during development.

Under this assumption, we first extracted the fundamental features of the branching pattern of the dorsal rami of the spinal nerves. The dorsal rami bifurcate into the medial and lateral branches, which innervate the multifidus and spinalis muscles, and the longissimus and iliocostalis muscles, respectively (Standring, 2015). The two branches end as medial and lateral cutaneous branches, after innervating the muscles. From this branching pattern, we created a basic building unit for our model; one main branch sends muscle-innervating twigs to a parallel pair of muscles and ends as one terminal sensory branch (Figure 1).

More importantly, in the human body, the dorsal and ventral rami exhibit a fundamental difference in branching design. While adjacent ventral rami from the plexus, except from those in the thoracic region, merge, nearby dorsal rami maintain the segmental innervation pattern all along the axial levels. We selected the muscle-innervating twigs that diverge from the root segment of the ventral rami from all spinal levels. Most of these branches are the first-order branch (not counting the dural sensory twig), and are unnamed or described as “nerve to” in gross anatomy textbooks. Because these unnamed branches do not join the plexus formation, finding the first-order branch from the ventral rami is equivalent manipulation to selecting the nerve branches that also maintain the segmental nature from the ventral rami. According to our working hypothesis, we assigned these “segmental” branches from the ventral rami to the innervation of muscles in the primaxial compartment. The selected muscles are listed in Table 1, and are described in the next chapter in detail.

**Table 1.**
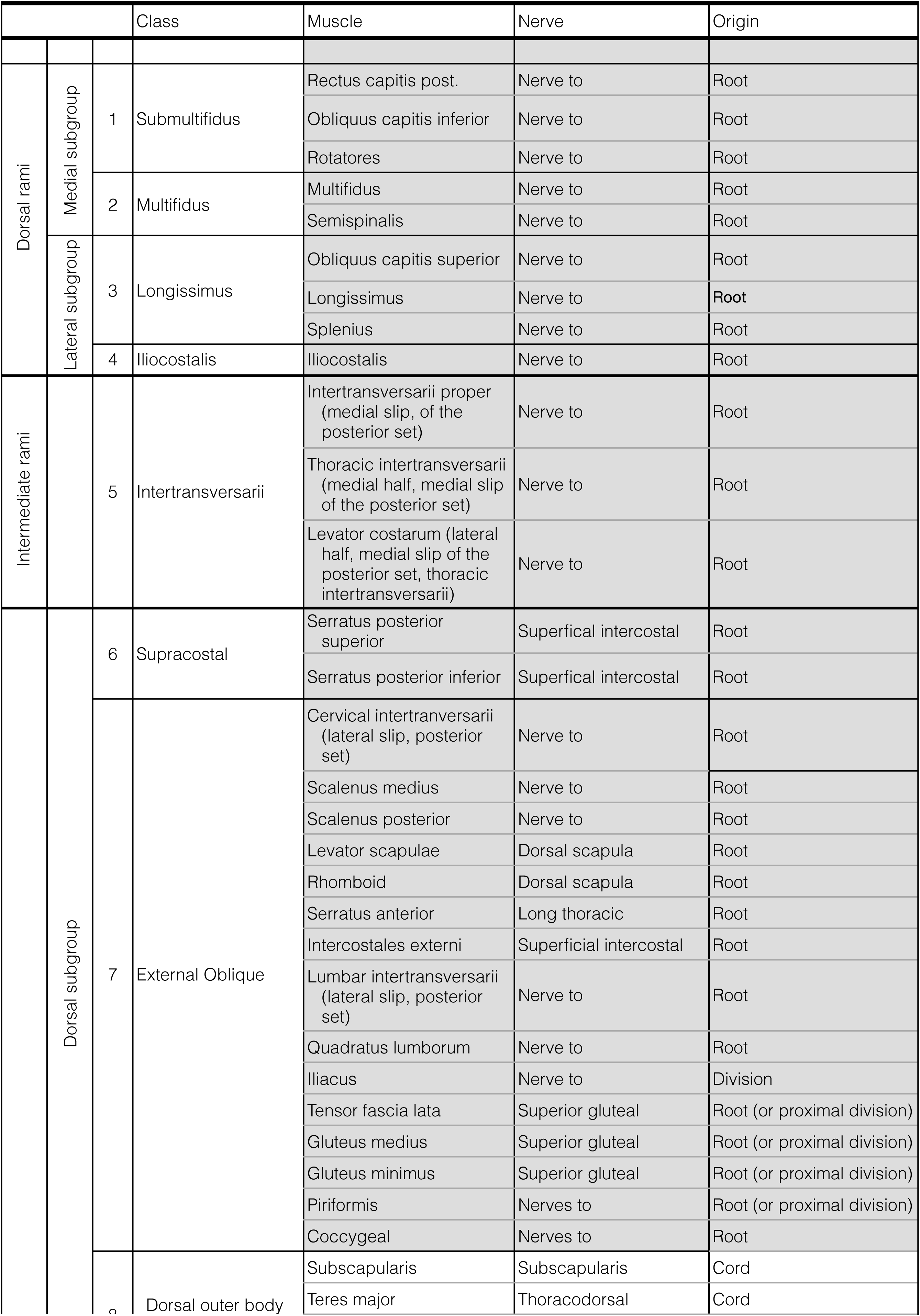
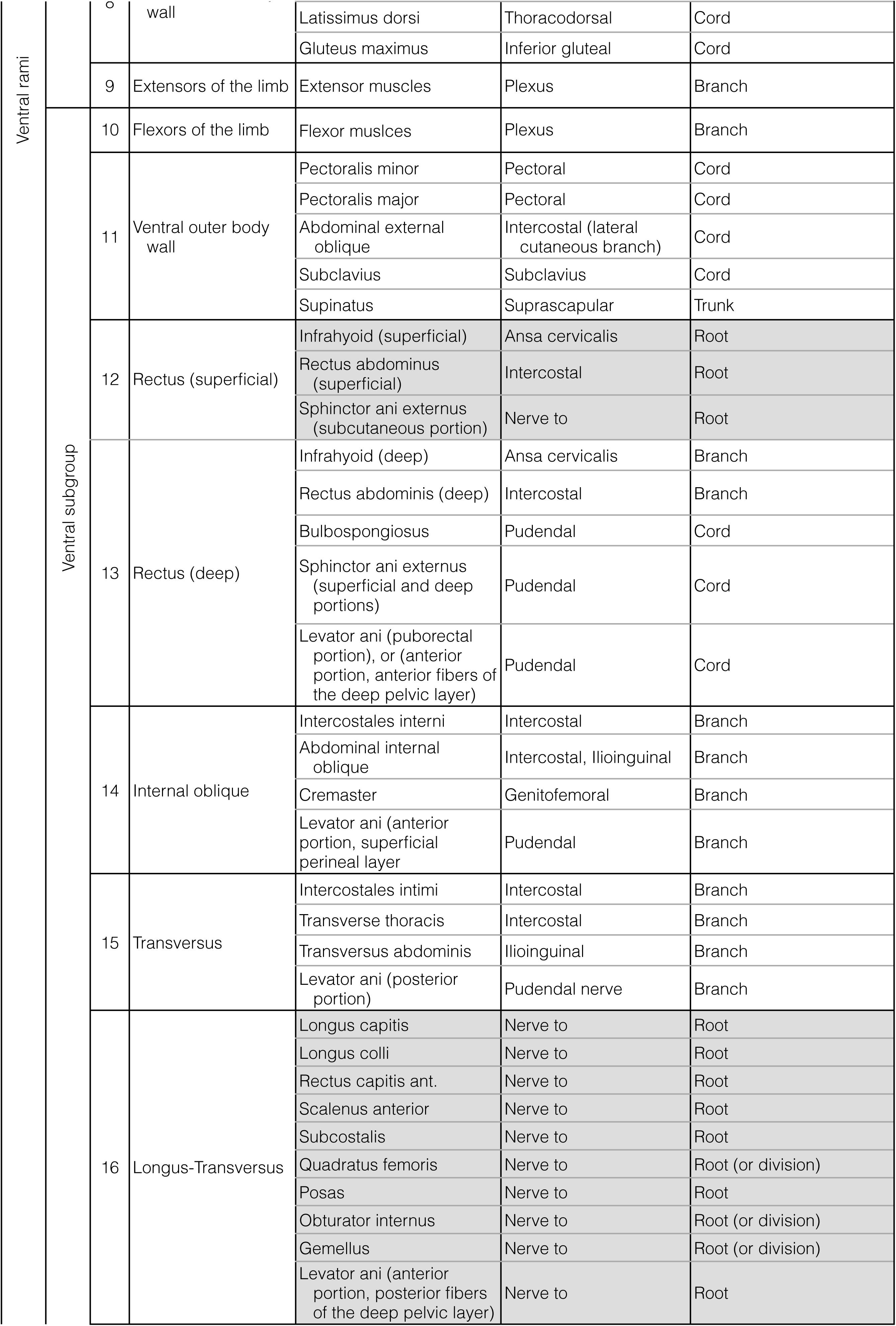
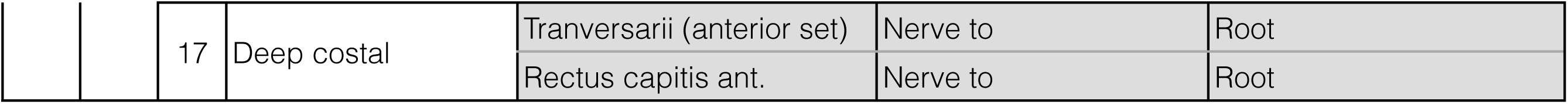
Classification of human muscles based on the view of the lateral somitic frontier.

We also needed another working hypothesis upon constructing a model. The second working hypothesis is that the homologous muscles at different axial levels of the body preserve the relative position to their parental motor nerve. Conversely, homologous spinal nerve branches maintain relative positions to the corresponding classes of surrounding muscles on their way to the targets. For example, we consider that the transverse abdominis and the innermost intercostal muscles are homologous because the intercostal nerve is positioned dorsally to them. Because of high variabilities in the functions and locations of muscles due to adaptive purpose, we cannot rely on these marks when we classify muscles into specific categories. Without this second hypothesis, we would lose critical landmarks for correspondence between a muscle and its innervation.

As muscles belong either to dorsal or ventral subgroups (Table 1; also see the next chapter), the primaxial compartment-responsible and outer body wall branches are also further subdivided to respective dorsal and ventral subbranches. The first-order branches for the primaxial compartment innervate the supracostal and external oblique muscle classes (dorsal subgroup) and the longus-transversus and deep costal classes (ventral subgroup; Figures 1 and 2). In addition to the above first-order branches to the primaxial muscles, our model adopted an additional primaxial compartment-responsible nerve branch that reaches the anterior midline, where it innervates muscles in the superficial rectus class (ventral subgroup), and further ends as another anterior primaxial cutaneous branch (Figures 1 and 2). We discuss in the next chapter about the reason for the introduction of the anterior primaxial branch into our model, in conjunction with the innervation to the rectus abdominis muscle, and explain that it plays significant roles in establishing the concept of dual innervation to the body wall by both primaxial and abaxial compartment-responsible anterior nerve branches.

Regarding the innervation of muscles in the abaxial compartment in the body wall, branches equivalent to the canonical intercostal nerves play the role of innervation for muscles in the internal oblique, transversus, and rectus (deep) classes. These abaxial-responsible branches also end as anterior cutaneous branches around the ventral midline region (classical anterior cutaneous branch of the intercostal nerve). This component of the branches is only seen in the ventral muscle subgroup; no dorsal counterpart exists for the dorsal muscle subgroup (Figures 1 and 2).

In line with the second working hypothesis, we emphasize that the primaxial compartment-responsible anterior branch (anterior primaxial branch) is not one of the collaterals from the canonical intercostal nerve, but is a distinctive entity because of the difference in trajectory in our model. The primaxial-responsible anterior branch changes its course to the superficial intermuscular space (e.g., between the external and inner intercostal muscles in the thoracic region) on its way to the anterior midline. In contrast, the canonical intercostal nerve maintains its course in the deep intermuscular space (e.g., between the inner and innermost intercostal muscles). Thus, our model highlights the two different anterior branches that reach the ventral midline and innervate two different classes (superficial and deep) of rectus muscles (Figures 1 and 2).

Another component, which is responsible for the abaxial muscle, is a novel class of nerve that runs together with the lateral cutaneous branch, namely the outer body wall branch (Figures 1 and 2). The outer body wall branch is such a unique motor nerve that it innervates the muscles that belong to the outer body wall and limb muscle classes, and also a sensory nerve that innervates the dermis in the abaxial compartment. On the other hand, the lateral cutaneous branch itself has conferred a unique anatomical status, which means that not only is it the cutaneous sensory nerve, but it also serves as a guideway to convey the outer body wall branch to the abaxial compartment.

The observations described below support the notion that the lateral cutaneous branch carries the motor and sensory innervation (outer body wall branches) to appendicular muscles and dermis. First of all, contrary to general perception, the lateral cutaneous nerve, but not the intercostal nerve, may be considered as a serial homolog to the nerve trunk in the brachial plexus because of the sensory innervation pattern to the axilla and medial arm regions. The equivalent nerve to the lateral cutaneous nerve from Th2, namely the intercostobrachial nerve, joins a branch from the brachial plexus to innervate the medial aspect of the arm (Standring, 2015; Loukas et al., 2006). This union indicates that the most posterior constituent of the brachial plexus is the lateral cutaneous nerve, as suggested by Cave (1929). Secondly, spinal nerves consisting of the brachial plexus also appear through the scalene gap, which is positioned on the anteriorly extended line of a series of points at which the lateral cutaneous nerve comes out of the skin in the trunk.

Furthermore, several previous reports have described the presence of a distinctive cutaneous branch that diverges from the lateral cutaneous branch, which has the original name of the extramural sensory branch (Koizumi and Horiguchi, 1992; Akita et al., 2002). However, we used the name of the outer body wall branch (outer branch in abbreviated form) in our model. The above studies demonstrated that the pectoral nerve from the brachial plexus further extends sensory terminal branches following the cognate motor innervation to the pectoral muscles. They also reported that distinctive nerve branches were diverged from the lateral cutaneous branches of the second to the fourth intercostal nerve, and ran on the fascial surface of the pectoralis major muscle with further traveling anteriorly to end as rudimentary sensory nerves to the lateral body wall. The reported sensory branch is different in trajectory such that the outer branch maintains extra dermic (on the fascial surface) path until it reaches a target, whereas the lateral cutaneous branch itself travels throughout the intra-dermis region.

The motor branch to the external oblique abdominis muscle takes a similar trajectory to those of the nerves described above. The nerve branch to the external oblique abdominis muscle diverges, shortly after the parental lateral cutaneous branch appears on the surface of the body wall, and then runs on the outer fascial surface to enter the muscles from its surface (Sato, 1973; Schlenz et al., 1999). On the other hand, direct twigs innervate the internal oblique abdominis and traverse abdominis muscles from the intercostal nerve itself that runs in the deep intermuscular space. As such, there are two different innervation patterns to the muscles associated with the body wall: direct innervation by twigs from the intercostal nerve for muscles within the body wall, and indirect innervation via the lateral cutaneous branch for those located on the surface of the body wall.

Under our working hypothesis, the outer body wall branch is different from the lateral cutaneous branches because they take different positions relative to the body surface muscles. Furthermore, our building unit assumes a novel class of the cutaneous branch to the tissues external to the body wall (Figures 1 and 2), and the reported sensory branches fulfill the requirements for the outer body wall branch in our model. These two notions led us to place a series of distinctive motor and sensory branches (outer body wall branches) that are accompanied by the lateral cutaneous branch all along the body axis in our model. The outer body wall branch also corresponds to the dorsal and ventral muscle subgroups by splitting into subbranches, which presumably run with the posterior and anterior subbranches of the lateral cutaneous branch, respectively.

Collectively, three components of nerve branches, which occupy different intermuscular spaces, differentially innervate muscles in the human body (Figures 1 and 2). The nerve branch that runs in the deep intermuscular space is a homolog to the canonical intercostal nerve and innervates body wall muscles in the abaxial compartment, terminating as an anterior abaxial cutaneous branch. The nerve branches that radiate from the root segment innervate primaxial muscles mainly located around the paraxial region, and one of them travels anteriorly in the superficial intermuscular space towards the ventral midline to innervate the primaxial muscle in the ventral body wall and ends as an anterior primaxial cutaneous branch. The outer body wall branch runs along the lateral cutaneous branch to innervate the dermis in the surface body wall and appendicular muscles.

The nature of the building unit in our model also presumes seven essential cutaneous innervations from the ventral rami all along the body axis (Figures 1 and 2). From dorsal to ventral in the ramification, the seven cutaneous branches are: 1) the branch from the dorsal division of the primaxial compartment-responsible nerve (posterior primaxial cutaneous branch); 2) the dorsal outer body wall cutaneous branch; 3) the posterior branch of the lateral cutaneous branch; 4) the anterior branch of the lateral cutaneous branch; 5) the ventral outer body wall cutaneous branch; 6) the anterior cutaneous branch of the canonical intercostal nerve (anterior abaxial cutaneous bench); and 7) the anterior cutaneous branch of the ventral division of the primaxial compartment-responsible nerve (anterior primaxial cutaneous branch). The number of cutaneous branches in our model is far higher than that previously described in the thoracic region, which is three (the anterior and posterior branches of the lateral cutaneous branch, and the anterior cutaneous branch of the intercostal nerve). The distribution of the seven cutaneous branches is described in detail when applying our model to the human body.

### 2. Muscle classes

Along with creating a model for the nerve branching pattern, we classified all muscles in the human body into seventeen categories under our third assumption that an equivalent class of muscles is innervated by a corresponding component of the spinal nerve, despite the various names of nerve branches at different axial levels. We based our classification according to Nishi (1938, see also Figure 6 in Sato, 1968), but with extensive modifications. Table 1 shows individual members in muscle classes. The proposed muscle classes are 1) submultifidus, 2) multifidus, 3) longissimus, 4) iliocostalis, 5) intertransversarii, 6) supracostal, 7) external oblique, 8) dorsal outer body wall, 9) extensors of the limb, 10) flexors of the limb, 11) ventral outer body wall, 12) superficial rectus, 13) deep rectus, 14) internal oblique, 15) transversus, 16) longus-transversus, and 17) deep costal.

#### 2 - 1) Back muscles and dorsal rami

The 1) submultifidus, 2) multifidus, 3) longissimus, and 4) iliocostalis muscle classes comprise the so-called intrinsic back muscles (primaxial muscle). The four muscle classes are subgrouped into medial (submultifidus and multifidus classes) and lateral subgroups (longissimus and iliocostalis classes). All muscles are innervated exclusively by the dorsal rami of the spinal nerve.

#### 2 - 2) Intertransversarii and intermediate rami

Multiple muscle slings between the transverse processes of the vertebrae comprise the intertransversarii muscle (Standring, 2015). The muscle complex of the intertransversarii consists of the anterior and posterior sets, between which the ventral rami of the spinal nerves run. The anterior set receives the innervation from the ventral rami of the spinal nerve, and thus belongs to the deep costal muscle class in our model (Sato, 1968; Standring, 2015). The posterior set further differentiates into the medial and lateral slips. The lateral slip also receives innervation from the ventral rami and is more related to the external intercostal muscle (Sato, 1968; Standring, 2015). Accordingly, we placed the lateral slip of the posterior set in the external oblique class. The medial slip of the posterior set corresponds to the intertransversarii muscle (Sato, 1968; Standring, 2015). Sato (1974) also found that the nerve branches to the actual intertransversarii muscle originated from the nerve root between the dorsal and ventral rami, and argued that they should therefore have a unique name; the intermediate rami. In agreement with Sato’s argument regarding the innervating rami, we placed the medial slip of the posterior set in the intertransversarii muscle class in the primaxial compartment (Table 1). In the thoracic region, the lateral half of the medial slip of the posterior set corresponds to the levatores costarum muscles (Sato, 1971 and 1974).

#### 2 - 3) Body wall and appendicular muscles, and ventral rami

The ventral rami of the spinal nerve innervate the muscles in the remaining twelve classes. We further separated them into either dorsal or ventral subgroups, based on the relative positions to the ventral ramus of the spinal nerve (Table 1). For example, in the case of the scalene muscles, the longus-transversus class, which includes the anterior scalene muscle, belongs to the ventral subgroup. In contrast, the external oblique class, containing the middle and posterior scalene muscles, belongs to the dorsal subgroup. In the case of appendicular muscles, the dorsal group includes the extensors, and the ventral group contains the flexor muscles.

The primaxial muscle classes in the body wall consist of the supracostal, external oblique, superficial rectus, longus-transversus, and deep costal classes. Although the branches from the ventral rami innervate muscles in the primaxial compartment, some muscles, especially those located close to the vertebrae, receive innervation from both dorsal and ventral rami. The branches from the dorsal rami sometimes innervate the rhomboid in addition to the dorsal scapular nerve (Aizawa and Kumaki, 1996; Saberi et al., 2017). Such double innervation has long been a matter of discussion in muscle classification. However, the dichotomy based on the dorsal-ventral rami is not suitable to use as a demarcation for primaxial-abaxial dichotomy. Instead, the muscles in the primaxial compartment receive such innervation by branches that they are segmental and diverge at the root segment of the spinal nerve as primary branches (Table 1).

What is unique in our model is that there exists the rectus muscle, which belongs to the primaxial compartments. Thus, contrary to general perception, we postulate the enclave of the primaxial compartment, even in the anterior body wall, which is supposed to belong exclusively to the abaxial compartment. We discuss the rationale for this in the next chapter.

Abaxial muscle classes include the dorsal outer body wall, extensors of the limb, flexors of the limb, ventral outer body wall, deep rectus, internal oblique, and transversus (Table 1). Limb and outer body wall muscle classes receive motor innervation (outer body wall nerve), which is conveyed by the lateral cutaneous nerve. The muscles in the internal oblique, transversus, and deep rectus classes are innervated by a homologous nerve to the intercostal nerve. These three classes of abaxial muscles belong to the ventral subgroup (Figures 1 and 2).

#### 2 - 4) Muscles in the pelvic floor

Compared to the definite muscular organization of the pelvic wall, that of the pelvic floor is inconsistent among standard textbooks and references, which resulted in significant confusion in terms of denoting individual muscles (Kearney et al., 2004). This difficulty may be attributed to the significant divergence of the muscular organization described in recent dissection studies from those in the standard reference.

In Gray’s Anatomy (Standring, 2015), for example, one of the muscular complexes in the pelvic floor, the levator ani muscle, consists of three muscle slings; the iliococcygeus, pubococcygeus, and puborectalis. The conception for the three parts in the levator ani muscle dates back to a study by Thompson (1899). However, later studies have revealed different landscapes on the muscular organization of the levator ani. Ayoub (1979) reported that the levator ani muscle consists of anterior and posterior parts, and that three layers exist in the anterior part. Bustami (1989) also reported a layered organization in the anterior portion of the levator ani muscle. In essence, the anterior portion is thick and further subdivided into deep and superficial layers, although the posterior portion is thin and aponeurotic. In the report, Bustami pointed out that the anterior fibers in the anterior portion (both superficial perineal and deep pelvic layers) correspond to the classical levator prostate (a subdivision of the pubococcygeus). In contrast, the posterior fibers in the superficial perineal layer of the anterior portion do not match any known named muscles, and the posterior fiber in the deep pelvic layer of the anterior portion corresponds to the classical pubococcygeus.

A recent 3-D reconstruction study also demonstrated that the levator ani muscle consisted of anterior and posterior portions, and that the single anterior head bifurcated into two discrete muscles in the posterior portion (Wu et al., 2015). Two muscle groups in the posterior portion of the levator ani muscle correspond to the classical puborectalis, and pubococcygeal and iliococcygeal muscles, which are collectively named pubovisceral muscles in that study. However, the pubovisceral muscle cannot physically be subdivided into pubococcygeal and iliococcygeal muscles. Instead, it is a bilayered muscle, consisting of medial and lateral layers. Furthermore, in Wu et al.’s study, the classical puborectalis muscle corresponds to the deep part of the external anal sphincter muscle.

The above three studies provide us with a completely different conception on the levator ani muscle. All studies agree the following two points: 1) the levator ani muscle is multi-layered; and 2) the classical puborectalis muscle does not correspond to any muscle fibers that were dissected. It was previously reported that the puborectalis part of the levator ani muscle is morphologically and functionally related to the external anal sphincter muscle (Shafik, 1975). Later, anatomical studies on the puborectalis muscle also argued the same point (Fröhlich et al., 1997; Fritsch et al., 2002).

The muscle organization of the external anal sphincter also possesses a similar conflict. A long-accepted view on the external anal sphincter is that it consists of three muscle components, superficial cutaneous, superficial, and deep components (Standring, 2015). In addition, Oh and Kark (1972) included another component of the levator ani muscle, the puborectalis muscle, into the external anal sphincter complex, for the apparent continuation of the deep component of the external anal sphincter to the puborectalis muscle, taking the number of components to four. Al-Ali et al. (2009) then reduced the previously recognized multiple parts to only two parts; the superficial cutaneous and deep parts of the external anal sphincter. The superficial part of the external anal sphincter includes only the superficial cutaneous muscle, and the deep part includes the classical superficial and deep muscles. Wu et al. (2015) also divided the external anal sphincter into two parts; the puborectal part, which includes the puborectalis, deep and superficial muscles of the external anal sphincter, and the cutaneous part. Nevertheless, all studies agree that the superficial cutaneous part of the external anal sphincter is an independent muscle component.

Although the number of subdivisions of the external anal sphincter varies depending on the study, the three components model of the external anal sphincter is useful when focusing on their continuities with neighboring muscles such as the perineal and levator ani muscles. Close examinations of the perineal regions have revealed that the deep component of the external anal sphincter was continuous with the puborectal part of the levator ani muscle, the superficial component of the external anal sphincter to the perineal muscles (Arakawa et al., 2010; Plochocki et al., 2016). Moreover, Ayoub (1979) also reported that the pelvic and middle layers of the levator ani muscle continue with the classical deep part of the external anal sphincter, and the perineal layer is continuous with the classical superficial part of the external anal sphincter, respectively. Baramee et al. (2019) also reported the integral architecture of the pelvic floor, which consists of the levator ani, perineal, and external anal sphincter muscles.

Collectively, the above dissection and 3-D reconstruction studies revealed a novel conception of the muscular organization in the pelvic floor, the muscles of which form a multi-layered structure and are continuous from the perineum to the anorectal regions. However, the muscle sheets differentiate into the morphologically segmented and functionally specialized complexes, including the perineal, external anal sphincter, and levator ani muscles, possibly because the muscles in the pelvic floor are responsible for highly specific functions such as micturition, ejaculation, and defecation.

Questions have arisen as to the reason why there are various descriptions about the anatomical organization of the pelvic floor, and the naming of the muscles. As described above, the external anal sphincter and levator ani muscles consist of multiple components. However, it is not clear whether the individual components are subdivisions of a single muscle mass or assembling elements for a building complex composed from different classes of muscles. If the latter is the case, it is challenging to compare the subdivisions of muscles in various studies without adequate reference categories. For this reason, we first distributed pelvic floor muscles into our muscle categories based on the stratification patterns of muscle components and continuities with neighboring muscles (Table 2).

**Table 2.** Allocation of pelvic floor muscles into the muscle classes in our model

|  |  |  |  |  |  |  |  |  |
| --- | --- | --- | --- | --- | --- | --- | --- | --- |
| Wu et al., 2015 |  | Deep transverse perineal | Superficial transverse perineal<br>Ischiocavernosum | Bulbospongiosum |  |  |  | Perineal muscle |
| Plochocki et al., 2016 |  |  | Superficial transverse perineal<br>Ischiocavernosum | Bulbospongiosum |  |  |  |  |
| Baramée et al., 2020 |  |  | Superficial transverse perineal<br>Ischiocavernosum | Bulbospongiosum |  |  |  |  |
| Oh and Kark, 1972 |  |  |  | Deep EAS<br>Puborectalis | Superficial EAS | Subcutaneous EAS |  | External anal sphincter (EAS) muscle |
| Al-Ali et al., 2009 |  |  |  | Upper part |  | Lower part |  |  |
| Wu et al., 2015 |  |  |  | Deep puborectalis | Superficial puborectalis | EAS proper |  |  |
| Plochocki et al., 2016 |  |  |  | Deep EAS | Superficial EAS | Subcutaneous EAS |  |  |
|  | **Longus-transverse** | **Transverse** | **Internal oblique** | **Deep rectus 2** | **Deep rectus 1** | **Superficial rectus 2** | **Superficial rectus 1** | **Muscle class** |
| Ayoub, 1979 |  |  | Anterior pelvic layer | Anterior middle layer | Anterior perineal layer |  |  |  |
| Bustami, 1989 | Anterior portion,<br>posterior fibers of the<br>deep pelvic layer | Posterior portion | Anterior portion,<br>superficial perineal<br>layer | Anterior portion,<br>anterior fibers<br>of the deep<br>pelvic layer |  |  |  | Levator<br>ani<br>muscle |
| Wu et al., 2015 |  | Medial pubovicelaris | Lateral pubovicelaris |  |  |  |  |  |
| Classic |  | Iliococcygeus | Pubococcygeus | Puborectalis |  |  |  |  |

Because the anorectal canal opens in the ventral midline of the body wall, we hypothesized that the surrounding muscle, the external anal sphincter, belongs to the rectus class of the muscles, and started with working the allocation of this muscle. In our model, all rectus muscles belong to one of the four subclasses (a pair in the primaxial rectus and a pair in the abaxial rectus). The standard three subdivisions of the external anal sphincter muscle are the subcutaneous part, superficial part, and deep part, which correspond to the superficial rectus and a pair of the deep rectus classes, respectively (Table 2).

We interpreted the continuations among pelvic muscles as the muscular integration of the body wall as the internal oblique abdominis muscle fascia connected with the rectus sheath that forms the complete circle of the abdominal wall. Because of the continuations of the bulbospongiosus muscle with the superficial external anal sphincter, we placed the bulbospongiosus muscle in the rectus muscle class (Plochocki et al., 2016; Table 2). Based on the fiber orientations, the other perineal muscles, the ischiocavernosus and transverse perineal muscles, were allocated to the internal oblique and transversus classes (Table 2). A significant part of the levator ani muscle belongs to the internal oblique and transverse muscle classes. However, a small part of the levator ani muscle, to which Bustami (1989) stated that no established subdivision of the levator ani corresponded, belongs to the longus-transversus class of muscle in the primaxial compartment (Table 2).

#### 2 - 5) Comparison with a mapping study

A previous mapping study in mouse demonstrated that muscles in the external oblique class, such as the rhomboids, levator scapulae, quadratus lumborum, levator costarum, and longus colli, belong to the abaxial compartment (Durland et al., 2008). This mapping is consistent with our classification; however, the status of some muscles from their study opposes our classification. They reported that the latissimus dorsi and rhomboids are mixed-muscle; the proximal portion of the intercostal muscles being primaxial, and the distal portion of those being abaxial. However, in our model, the latissimus dorsi entirely belongs to the abaxial compartment, and all of the rhomboids belong to the primaxial compartment. Regarding the intercostal muscles, the external intercostal muscle is in the primaxial compartment, whereas the innermost and internal intercostal muscles are in the abaxial compartment. Therefore, the difference in species between human and other mammals may explain the inconsistency. We, however, did not rule out the possibility that some muscles fit in different compartments from those predicted in our model, because our classification is highly schematic and hypothetical. Nevertheless, the previous mapping is mostly in agreement with our muscle classification.

### 3. Detailed application of our model to the human body

We deployed our model in the peripheral innervation pattern of the human body and depicted it below in detail. Our illustrations on the innervation pattern include anatomical evidence and provide a great deal of practical information, although we understand that these are hypothetical.

#### 3 - 1) Thoracic region

We begin with the description of the peripheral innervation pattern with the thoracic region first, because of its relative simplicity. The intercostal nerves radiate first-order branches from the root portion to the ventral subgroup of primaxial muscles, such as the subcostalis in the longus-tranversus class, and the anterior set of the intertransversarii muscle in the deep costal class (Figure 3 and Table 1).

**Figure 3.**
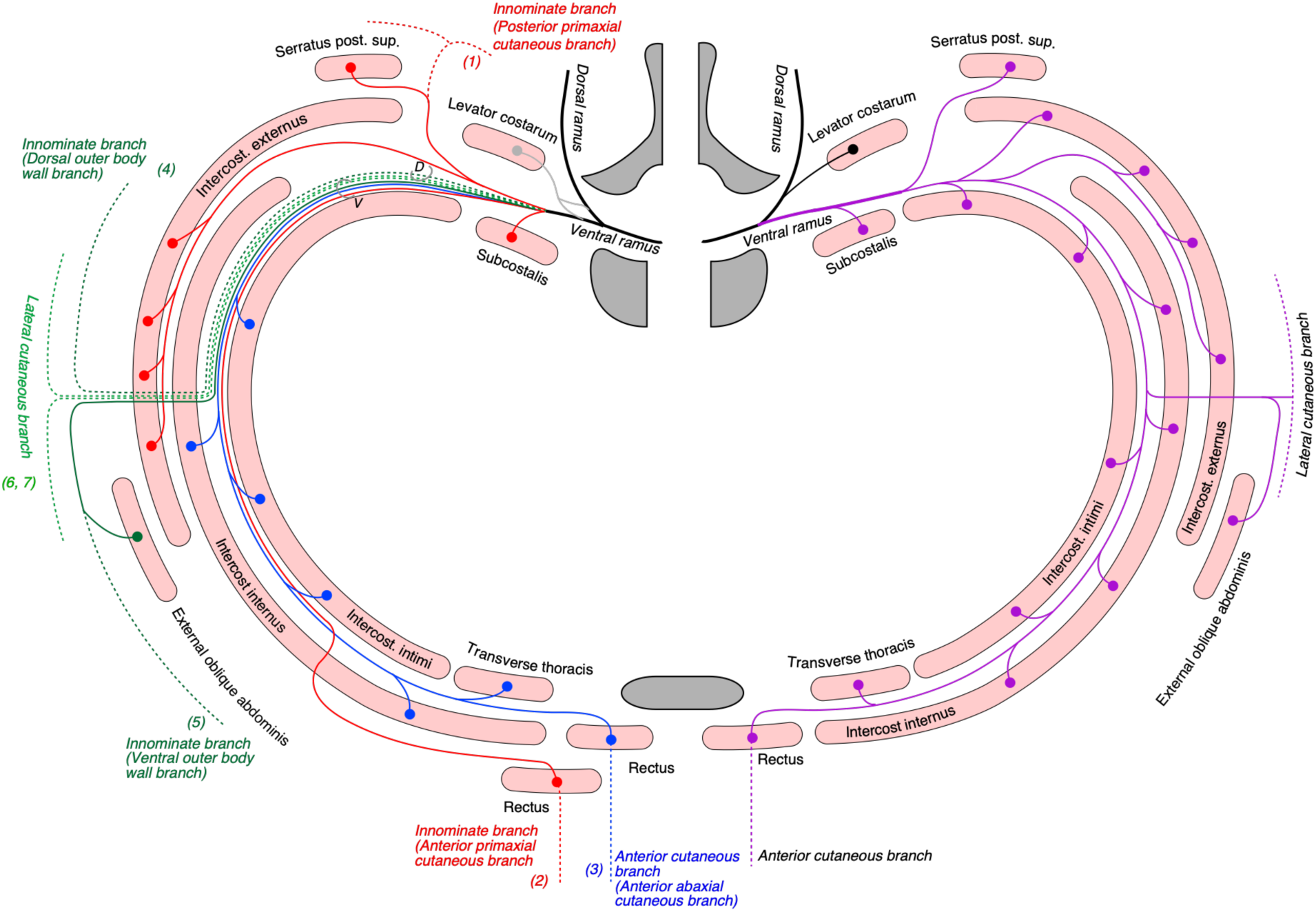
Hypothetical innervation pattern of three nerve components of the intercostal nerve in the mid-thoracic level. Because muscles and nerves are projected onto a single transverse plane (around Th5–6), not all of them are necessarily localized in the same transverse section, as illustrated. The schematic drawing in the right half of the figure shows the canonical branching pattern of the intercostal nerve, as is generally perceived. The single intercostal nerve (purple line) sequentially innervates muscles on its way to the rectus muscle. The branching pattern in the left half is based on our model. The blue lines are the canonical intercostal nerve and muscle twigs. The red lines are the nerve branches responsible for the primaxial compartment. The dark and light green lines are for the outer body wall and lateral cutaneous branches, respectively. The vertebra (upside) and sternum (downside) are dark gray in color. The numbers in parentheses indicate the seven cutaneous branches, which are presumed to be present in our model. Nerve branches that innervate muscle in the dorsal and ventral subgroups are in bundles of (D) and (V), respectively.

Regarding the first-order branches to the dorsal subgroup of primaxial muscles from the intercostal nerve, a problem arose when we tried to find a segmental branch, because such a specific branch had not yet been recognized. However, we found an interesting report that a specific muscle group, including the external intercostal muscle and some supracostal muscles such as the serratus posterior muscle, are innervated by an independent branch from the root of the intercostal nerve (Kodama, 1986). Although this branch, namely the superficial intercostal nerve, has not been adopted in the Terminologica Anatomica and is not described in gross anatomy textbooks, Kodama argued for the significance of this branch for following reasons.

Standard textbooks of human gross anatomy simply describe that the external intercostal muscle receives innervation from the intercostal nerve. However, it is equivocal how the intercostal nerve gains access to the external intercostal muscle. The intercostal nerve sends innervating twigs, which pass through the internal intercostal muscle, or sprout collateral into the intermuscular space between the external and internal intercostal muscles. However, according to Kodama (1986), the superficial intercostal nerve diverges from the root segment, travels between the internal and external intercostal muscles, and innervates only the external intercostal muscle. The intercostal nerve itself innervates only the internal and innermost intercostal muscles, but never the external intercostal muscle. Moreover, the superficial intercostal nerves maintain their trajectories under the lower margin of the costa in the lower thoracic region, although the main intercostal nerve (thoracolumbar nerve) deviates from the costa, traveling towards the abdominal region. Kodama argued that these two differences in nerve trajectories are enough to grant unique status to this unfamiliar nerve branch. He also found that the superficial intercostal nerve sometimes reached the anterior abdomen and innervated the rectus muscle (see below).

The superficial trajectory and divergence from the root segment of the intercostal nerve are relevant to our criteria of the primaxial muscle-responsible branch. We adopted the superficial intercostal nerve as a segmental component of the ventral rami to the dorsal subgroup of primaxial muscles. Thus, the nerve innervates the dorsal subgroup of the muscles in the primaxial compartment, including the serratus posterior (supracostal class) and the external intercostal muscle (external oblique class; Figure 3).

The canonical intercostal nerve innervates abaxial muscles, such as the internal and innermost intercostal muscles (internal oblique and transverse classes) and the rectus abdominis (deep rectus class; Figure 3).

The external oblique abdominis muscle is one that belongs to the ventral outer body wall class, and receives a motor branch from the lateral cutaneous branch in the lower thoracic region (Sato, 1973; Sakamoto et al., 1996; Schlenz et al., 1999; Figure 3). No muscle in the dorsal outer body wall class is in the thoracic region.

The general perception of the cutaneous sensory innervation on the thoracic wall is that the lateral cutaneous branch plays a sensory role for the lateral portion, together with the anterior cutaneous branch for the anterior portion. However, our model hypothesizes that three more cutaneous branches exist on the thoracic wall; the dorsal and ventral outer body wall branches, and the primaxial anterior branch. Further study is necessary to confirm whether the predicted cutaneous branches are present or not. However, some cutaneous branches with predicted innervation patterns have been reported (Aizawa and Kumaki, 1996).

#### 3 - 2) Rectus muscle

In our model, two anterior branches exist in a single intercostal space (Figures 1– 3). In essence, one branch, which innervates putative primaxial muscles, travels in the superficial intermuscular space (anterior primaxial branch), and the other corresponds to the canonical intercostal nerve, runs in the deep intermuscular space and innervates the putative abaxial muscles (anterior abaxial branch). We describe the reasons for introducing the concept of the dual innervation below.

In the lower thoracic and upper lumbar regions, we were puzzled by the relative position of the rectus muscle to the internal oblique abdominis muscle. Above the arcuate line, the posterior lamina of the internal oblique abdominis is adjoined to the posterior lamina of the rectus sheath. However, below the arcuate line, posterior lamina of the internal oblique abdominis muscle runs anterior to the rectus abdominis muscle, fusing with the anterior lamina of the rectus sheath. This laminae arrangement of the rectus sheath indicates that the position of the rectus abdominis is anterior to or on the same transverse plane as the internal oblique above the arcuate line. However, below the arcuate line, the rectus abdominis is positioned posteriorly to not only the internal oblique, but also the transverse abdominis muscle.

This topological difference in the rectus muscle may present two alternatives; the rectus abdominis passes through the internal oblique abdominis at the arcuate line, or two different subclasses, superficial and deep rectus muscles, are distributed above and below the acute line, respectively. We adopted the latter interpretation in our model, and further allocated the superficial rectus muscle to the primaxial one, which is innervated by the primaxial compartment-responsible nerve (anterior primaxial branch) and the deep rectus to the abaxial nerve, which is innervated by the canonical intercostal nerve (anterior abaxial anterior branch).

The rationale for this mapping is as follows. The superficial intercostal nerve mentioned above, which is responsible for the dorsal subgroup of muscles in the primaxial compartment, occasionally innervates the rectus abdominis muscle (Kodama, 1986). Because the rectus abdominis belongs to the ventral subgroup of muscles, it seems odd that the superficial intercostal nerve for the dorsal subgroup of muscles innervates the rectus abdominis, which belongs to the ventral subgroup. This mismatch necessitated us to introduce a nerve branch that is obligated to the ventral primaxial muscle (e.g., superficial rectus abdominis). Thus, the rectus abdominis receives either the primaxial-responsible anterior branch, or the canonical intercostal nerve, depending on the axial levels; however, the trajectories are different. Because of the superficial location of the primaxial rectus muscle, a cognate nerve branch diverges from the common trunk of the spinal nerve on the way to the anterior midline in order to change the trajectory to the superficial intermuscular space between the external and inner intercostal muscles in the thoracic region, or between the external and internal oblique abdominis muscles in the abdominal region. On the other hand, the canonical intercostal nerve maintains its deep trajectory between the inner and innermost intercostal muscles or between the internal oblique and transverse abdominis muscles. In the above case, where the rectus abdominis receives the innervation by the superficial intercostal nerve, we reasoned that the nerve branch to the superficial rectus might have changed its trajectory at the very proximal segment of the spinal nerve, and consequently merged into the superficial intercostal nerve.

After the innervation of the rectus abdominis muscles, both the canonical intercostal and primaxial-responsible nerves end as the anterior cutaneous branches of the spinal nerve. However, the locations of their points of exit may differ around the ventral midline, as shown in a standard gross anatomy atlas, which depicts two parallel series of exit points of the anterior cutaneous branches in the anterior thorax (Figure 253 in Netter, 2014).

#### 3 - 3) Lumbar region

In the upper lumbar region, the primaxial muscles in the dorsal subgroup, including the quadratus lumborum, iliacus, tensor fascia lata, and both the gluteus medius and minimus (external oblique class), and the psoas in the ventral subgroup (longus-transversus class), receive innervation by the primaxial-specific branches, which are unnamed or described as “nerve to” (Figure 4). However, the branch to the gluteus muscles of the dorsal primaxial muscle-responsible nerve is called the superior gluteal nerve. Moreover, the terminal cutaneous branch of the superior gluteal nerve has a sensory extension (Figure 4; Akita et al., 1992).

**Figure 4.**
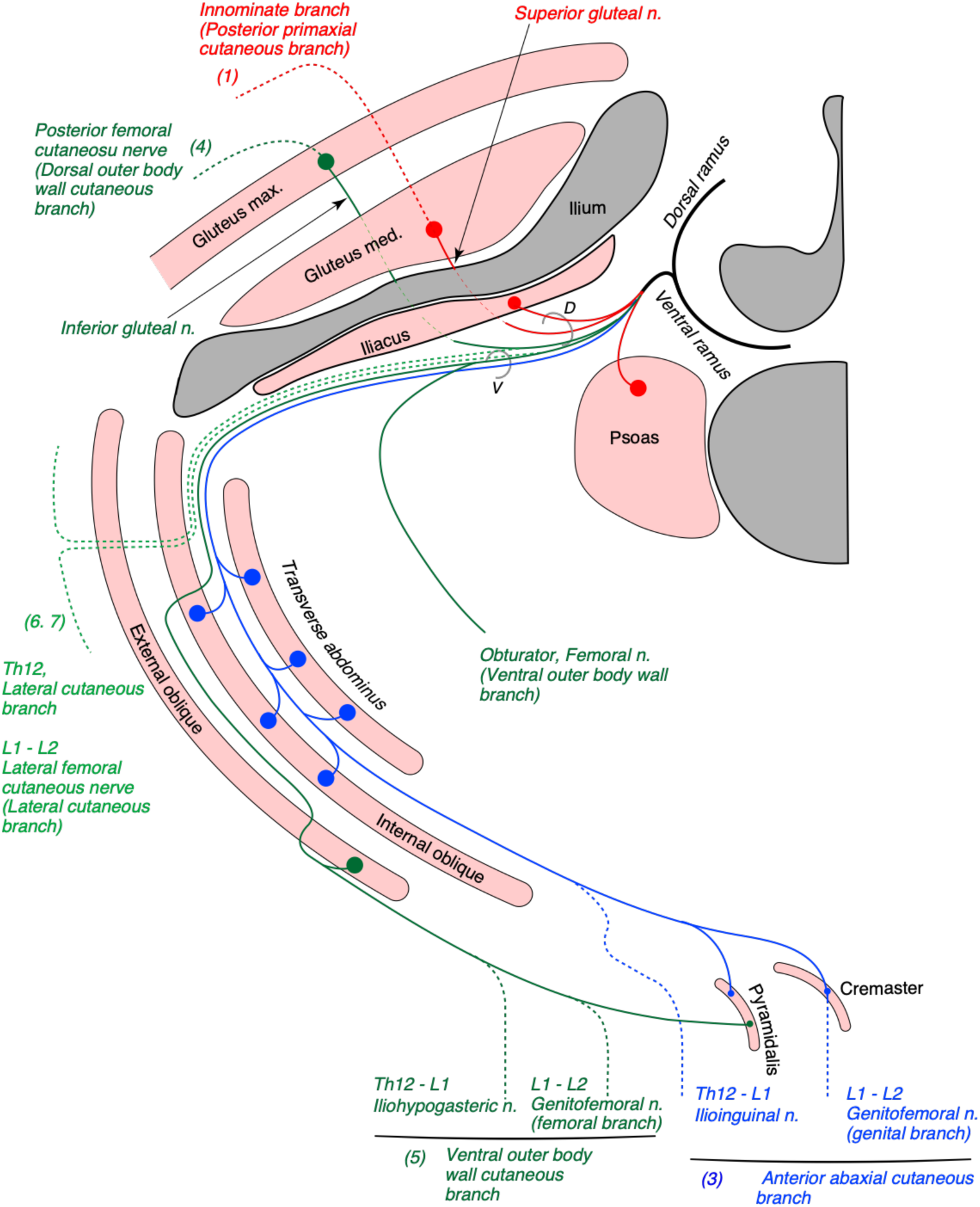
Hypothetical innervation pattern in the upper lumbar region. Nerve branches are super-imposed on a hypothetical single plane; therefore, not all of them necessarily exist on the same transverse plane. The named nerve branches are as indicated. Otherwise, they are innominate. The blue lines are the lumbar counterpart to the canonical intercostal nerve (the ilioinguinal nerve and genital branch of the genitofemoral nerve). The red lines are the nerve branches responsible for the primaxial compartment. The dark and light green lines are the outer body wall nerves, and the lateral cutaneous branches (the lateral femoral cutaneous nerve), respectively. The numbers in parentheses indicate the seven cutaneous branches (hatched segments) in our model. The axial origins of the nerve branches in the upper lumbar plexus are indicated for clarity. The nerve branches that innervate muscle in the dorsal and ventral subgroups are in bundles of (D) and (V), respectively. Bony structures are shaded in dark gray.

The spinal nerve from the lumbar spinal segment 1 (L1) usually bifurcates into the iliohypogastric and ilioinguinal nerves (Standring, 2015). In our model, the iliohypogastric nerve is equivalent to the outer body wall nerve. It conveys motor innervation to the external oblique abdominis muscle and sensory fiber to the anterolateral portion of the lower abdomen. Contrary to the general description in standard textbooks, the iliohypogastric nerve does not contain motor fibers to the transverse and internal oblique abdominis muscles, because these two muscles belong to different muscle classes, such as the internal oblique and transverse classes. On the other hand, it is the ilioinguinal nerve that is homologous to the canonical intercostal nerve, and conveys motor innervation to the internal oblique and transverse abdominis muscles. The ilioinguinal nerve then ends as a sensory nerve to the skin around the genital area of the abaxial compartment. We emphasize that the iliohypogastric and ilioinguinal nerves take different trajectories. The iliohypogastric nerve changes its trajectory from the deep to the superficial intermuscular spaces around the lateral body region (around the superior anterior iliac spine). In contrast, the ilioinguinal nerve maintains the deep intermuscular trajectory to a more anterior portion of the body wall (until the superficial inguinal ring).

Standard textbooks describe that the genitofemoral nerve is the main trunk from the L2 axial level, and also bifurcates into the genital and femoral branches. In our model, the femoral branch is the homolog of the outer body wall nerve and carries cutaneous sensory innervation to the upper medial thigh. The genital branch is the homolog to the canonical intercostal nerve. It conveys motor innervation to the cremaster muscle (a derivative of the internal oblique muscle) and sensory innervation to the genital area. The separation of the genitofemoral nerve into different categories of nerve branches reflects their trajectories within the abdominal wall. The genital branch of the genitofemoral nerve takes a deep trajectory, as described in gross anatomy textbooks. The branch enters the intermuscular space through the deep inguinal canal, in which it continues forward, and appears out of the superficial inguinal canal. On the other hand, the femoral branch runs on the fascial surface of the psoas major in no association with the abdominal muscles, and appears toward the upper medial thigh behind the inguinal ligament (the aponeurosis of the external oblique muscle).

The muscles in the lower limb (appendicular muscle subclass) are innervated by the outer body wall branches (the femoral and obturator nerves) that run along with the lateral cutaneous branches from the third and fourth lumbar spinal nerves. The dorsal and ventral extensions of the outer body wall nerve are the posterior and lateral femoral cutaneous nerves, respectively. This assignment of the outer body wall nerve partly comes from the fact that the inferior gluteal nerve, which is the dorsal outer body wall branch in our model, sometimes sends out the posterior femoral cutaneous nerve (Nakanishi et al., 1976).

#### 3 - 4) An explanation for variable branching pattern of the lumbar plexus

The branches of the spinal nerves from the lumbar (L) 1 and 2 spinal levels, the iliohypogastric, ilioinguinal, and genitofemoral nerves, are highly variable in many aspects of innervation patterns (Al-dabbagh, 2002; Peschaud et al., 2006; Nidiaye et al., 2010; Cirocchi et al., 2019). These branches are often divided and communicate with each other at various regions of the anterolateral abdominal wall, and travel in the different intermuscular spaces, piercing the internal and external oblique abdominis muscles and appearing at various points of the aponeurosis of the external oblique. Moreover, some of the branches from L1 and 2 spinal nerves are sometimes missing. We describe below the logic of how highly mutable innervation patterns in the lower abdomen emerge from our hypothetical innervation pattern.

A previous detailed study demonstrated that the L1 and L2 spinal nerves always bifurcate in more or less degree into superficial and deep branches (Kumaki, 1981 and 1995; see also Tokita, 2006). The superficial branch first runs in the deep intermuscular space between the transverse and internal oblique abdominis muscles. However, it changes course to the superficial intermuscular space between the internal and external oblique abdominis muscles, comes out through the external oblique aponeurosis around the anterior superior iliac spine, and travels anteriorly to innervate the suprapubic skins. This superficial branch resembles the trajectory of the typical iliohypogastric nerve, as described in standard anatomy textbooks. The deep branch takes similar trajectories to the canonical intercostal nerve, running in the deep intermuscular space throughout its course to come out from the rectus abdominis muscle, and may represent the typical ilioinguinal nerve. There is a third class of branches with a mixed trajectory, which runs in the deep portion but does not enter the rectus abdominis, and instead emerges from the surface of the external oblique abdominis.

Kumaki also found that the thickness of the superficial and deep branches is reciprocal, depending on what axial levels of the spinal nerve are predominant to the formation of the lumbar plexus (Kumaki, 1981 and 1995). The superficial branch is conspicuous if the posterior spinal nerves mainly constitute the lumbar plexus. In contrast, the deep branch is evident when spinal nerves from more anterior axial levels contribute to the lumbar plexus formation (e.g., the contribution of Th12 to L1). The above-mixed type of branch also appears when the more anterior spinal nerves are also the primary composite. In an extreme case where the Th12 (or the subcostal nerve) and L1, but not L2, spinal nerves constitute the lumbar plexus, the superficial branch (iliohypogastric nerve) from the L1 spinal nerve and the genitofemoral nerve (femoral branch) from L2 are both smaller in diameter. In contrast, the deep branches (the ilioinguinal nerve and the genital branch of the genitofemoral nerve) are wider in diameter. The above study also reported that the lateral cutaneous branch is present from the lumbar spinal nerve (L1) only when the deep branch is noticeable. Otherwise, the lateral femoral cutaneous nerve appears from the lumbar spinal nerves.

Kumaki (1981 and 1985) argues that the anterior or posterior shift in the axial levels of the spinal nerves that contribute to the formation of the lumbar plexus generates the variable branching patterns. Furthermore, Kumaki also stated that the axial shifts of the plexus-forming spinal nerves may reflect the relative proportion of innervation targets, the pre-girdle (actual body trunk) and girdle (of the lower limb) areas within the lower abdominal wall. In our model, the superficial branch (outer body wall branch) is responsible for the innervation of muscles and dermis that are related to the appendicles (girdle) and the deep branch, which is equivalent to the canonical intercostal nerve, responsible for muscles and dermis in the body wall (pre-girdle; Figure 4). Consistent with our model, the genital branch of the genitofemoral nerve (equivalent to the intercostal nerve) innervates the dermis and cremaster muscle in the body wall, whereas the femoral branch (outer body wall nerve) innervates the dermis in the upper medial thigh. Thus, our model theorizes that axial fluctuation of the border between the pre-girdle (body) and girdle (appendicle) creates the various sizes and piercing points of the bifurcated branches from the L1 and L2 spinal nerves, consequently leading to the variable branching patterns of the lumbar spinal nerves in the lower abdomen.

#### 3 - 5) Body all organization

Innervation patterns in the thoracic and abdominal regions in our model may lead to radical changes of view on the muscle organization of the body wall (Figure 5). The muscles in the thoracic, abdominal, and pelvic walls have three layers (the cutaneous muscle layer is not counted because of its absence in human; Nishi, 1938, Hall et al., 2017). The traditional view on the homology of muscles between the thoracic and abdominal walls is that the external intercostal muscle corresponds to the external oblique abdominis, the intercostal interni to the internal oblique abdominis, and the intercostal intimi to the transverse abdominis. This perceived conception is based on the topology of muscles in the body wall, and partly comes from the running orientation of muscle fibers. However, our view, which is based on the primaxial-abaxial distinction, is that the external intercostal muscle is homologous to the quadrates lumborum (motor innervation specific to muscles in the primaxial compartment); the intercostal interni to the internal oblique abdominis; and the intercostal intimi is to the transverse abdominis (innervation by the canonical intercostal nerve homolog, abaxial compartment-responsible branch). The external oblique abdominis muscle is not homologous to the intercostal externi; instead, it is more closely related to muscles such as the latissmisus dorsi, because it belongs to the outer body wall muscle class (innervation via the lateral cutaneous branch).

**Figure 5.**
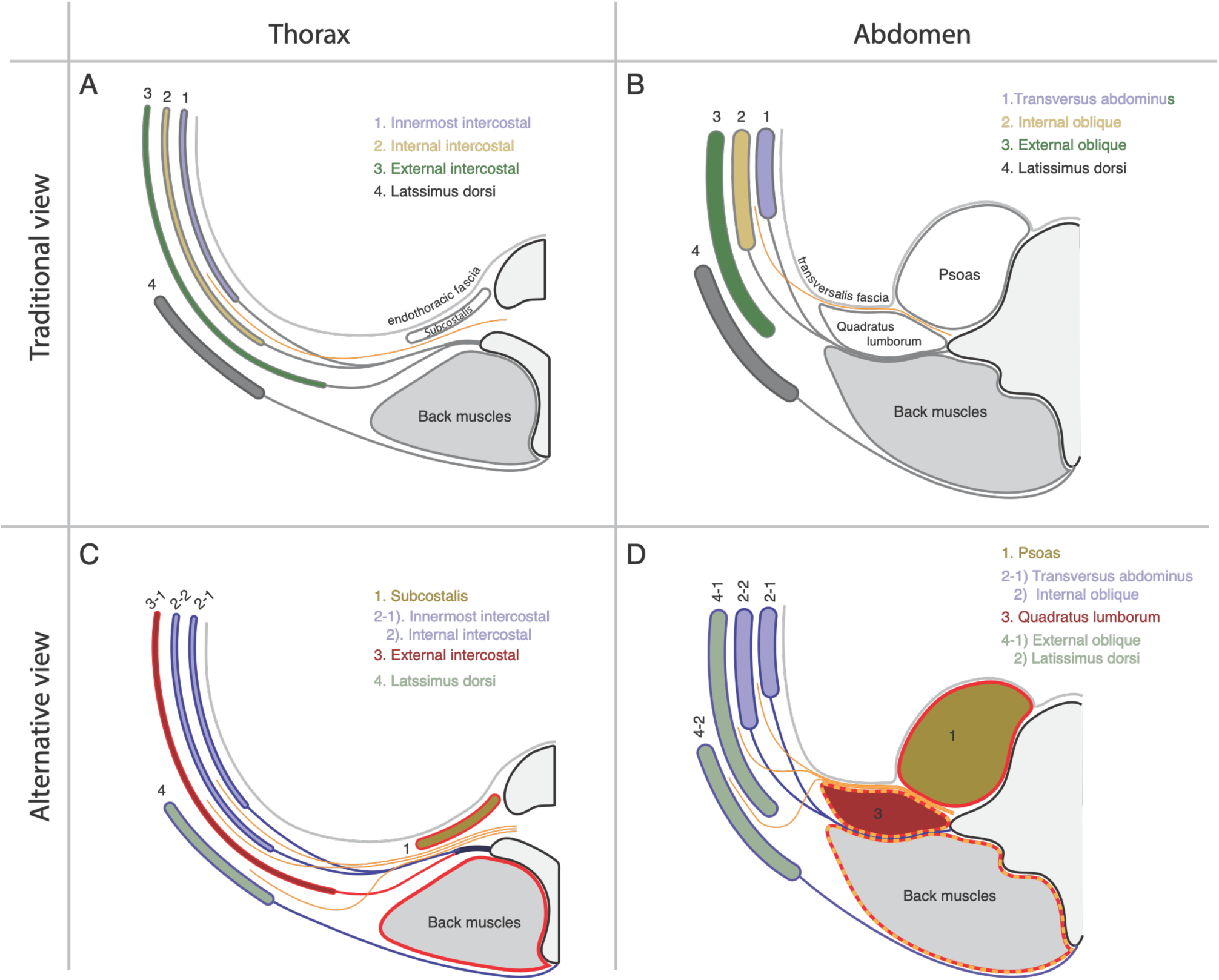
Layer organization in the body wall. The homology between the thorax (A and C) and abdominal wall (B and D) is color-coded. The traditional view is in the upper half (A and B). Here, the external intercostal muscle corresponds to the external oblique abdominis muscle, the internal intercostal muscle to the internal oblique abdominis muscle, and the innermost intercostal muscle to the transverse abdominis muscles. The lower half shows an alternative view from our model (C and D). The subcostalis muscle is equivalent to the psoas muscle, and the internal and innermost intercostal muscles correspond to the internal oblique and transverse abdominis muscles. The external intercostal muscle is homologous to the quadratus lumborum. The external oblique abdominis muscle is more closely related to the latissmus dorsi. In C and D, muscle-investing fasciae in the primaxial and abaxial compartments are shown in red and blue lines, respectively, and the thoracolumbar fascia is indicated by orange dashed lines. The spinal nerve is in yellow. In C and D, the three essential branches run in the different intermuscular spaces.

The above muscle homology in the body wall also strongly indicates that the primaxial and abaxial compartments are in different configurations between the thoracic and abdominal regions, hence the positions of the lateral somitic frontier. In the thoracic region, the abaxial compartment is juxtaposed with the primaxial compartment, whereas the abaxial compartment adjoins the primaxial compartment at the lateral raphe in the lumbar region (Figure 5).

The fascial organization in the lumbar region matches the body wall organization based on the primaxial-abaxial distinction to a certain extent (Figure 5). The thoracolumbar fascia invests the intrinsic back muscles and quadratus lumborum, and the external oblique abdominis muscle is located entirely external to the thoracolumbar fascia (Willard et al., 2012; Standring, 2015). Thus, the thoracolumbar fascia may represent an investing fascia to wrap the intrinsic back muscles and the dorsal group of muscles in the primaxial compartment. An adjoining point of the thoracolumbar fascia with the fascia of the transverse abdominis, the so-called the lateral raphe, has to date been interpreted as the demarcation of the epaxial and hypaxial compartments. In our interpretation, the primaxial compartment meets the abaxial compartment at the lateral raphe in the abdominal region. The psoas has an independent fascia from the thoracolumbar fascia, and is in the distinctive compartment, also termed the psoas compartment, but this is likely a result of the separation of the psoas fascia from the thoracolumbar fascia due to the presence of the spinal nerve between the psoas and quadratus lumborum. Indeed, the psoas-investing fascia lies on the anterior layer of the thoracolumbar fascia where the spinal nerve is absent, thus constituting a primaxial compartment-containing fascial system together with the thoracolumbar fascia.

#### 3 - 6) Pelvic region

Regarding muscles in the pelvic wall in our model, the dorsal group of muscles in the primaxial compartment includes the gluteus medius, gluteus minimus, piriformis, tensor fascia lata, piriformis, and gemellus (external oblique class), all of which are innervated by the superior gluteal nerve or direct branches from the sacral plexus, as previously reported in the literature (Figure 6). The ventral subgroup of muscles in the primaxial compartment includes the obturator internus and quadratus femoris (longus-transverse class). The direct branches from the sacral plexus innervate these muscles, as is also mentioned in textbooks (Figure 6).

**Figure 6.**
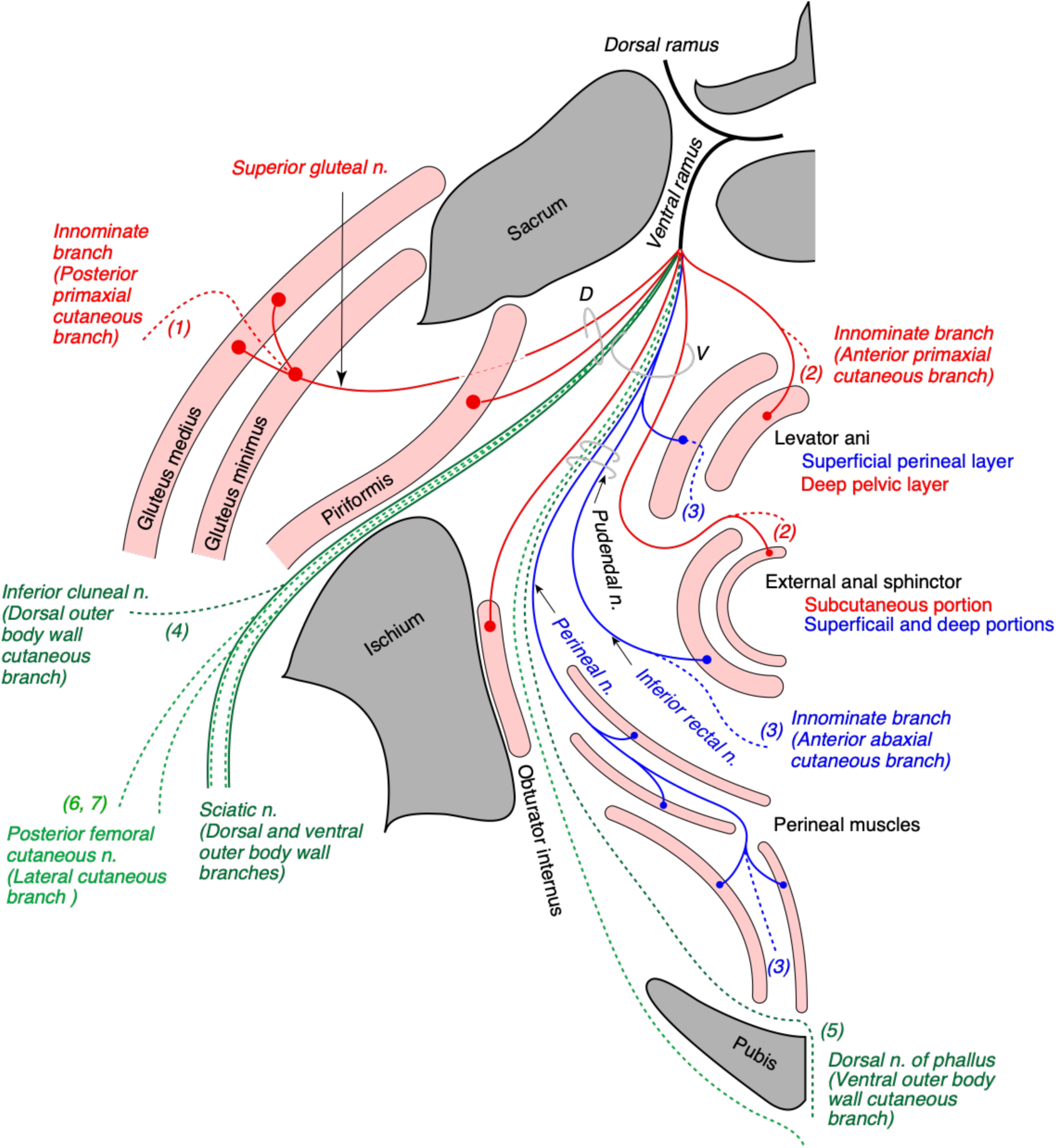
Schematic view of the innervation pattern of three main nerve components in the pelvis. The superior gluteal nerve (red) innervates the gluteus medius and minimus, and its cutaneous extension is innominate (1. posterior primaxial cutaneous branch). The levator ani muscle receives differential innervation of primaxial (red) and abaxial (blue) anterior branches; the perineal layer of the levator ani muscle and the deep pelvic layer are innervated by the abaxial and primaxial anterior branches, respectively. Similarly, the subcutaneous part of the external anal sphincter is innervated by the anterior primaxial branch, and the anterior abaxial branch (the inferior rectal nerve) is responsible for the rest of the external anal sphincter muscle. The muscles associated with the perineal membrane are innervated by the pudendal nerve (perineal nerve, blue). All anterior branches terminate as cutaneous branches (hatched lines), as marked as (2, anterior primaxial cutaneous branch) and (3, anterior abaxial cutaneous branch). The sciatic nerve is the outer body wall branch (dark green). Unlike in the brachial plexus, branches in the sacral plexus are not segregated into the dorsal and ventral divisions. The inferior cluneal nerve corresponds to the dorsal outer body wall nerve (4, dark green). The dorsal nerve of the phallus is a homolog to the ventral outer body cutaneous nerve (5, dark green). The posterior femoral cutaneous and posterior scrotum (labial) nerves are equivalent to the lateral cutaneous branch (6 and 7, light green).

Innervation of the muscles in the pelvic floor involves two trajectories from the sacral plexus. They receive the innervation via the pudendal nerve and direct branches from the sacral plexus. In Gray’s Anatomy (Standring, 2015), the primary nerve branch to the levator ani muscle diverges from the pudendal nerve and is often called the inferior rectal nerve. Also, the levator ani muscle receives its innervation directly from the sacral plexus, which is termed the levator ani nerve, and is often marked as a variant innervation from the inferior rectal nerve (Bustami, 1989; Grigorescu et al., 2008; Wallner et al., 2006 and 2008; Nyangoh Timoh et al., 2017). Bustami (1989) also reported the specific innervation of the direct sacral branch to a distinctive part of the levator ani muscle. Dual innervation to the external anal sphincter is also well documented; direct sacral branches predominantly innervate the subcutaneous part of the external anal sphincter, and the branches from the pudendal nerve innervate the rest portion of the external anal sphincter (Wallner et al., 2006 and 2008; Schraffordt et al., 2004; Plochocki et al., 2016).

Our model postulates two things: 1) the existence of the primaxial and abaxial parts of the rectus muscle; and 2) the differential innervation to the primaxial and abaxial rectus muscles by a direct branch from the sacral plexus and the pudendal nerve, respectively (Figure 6). Accordingly, the subcutaneous part of the external anal sphincter and a specific portion of the levator ani muscle, both of which are innervated by the direct branches from the sacral plexus, correspond to the primaxial rectus muscle. The other portions of the external anal sphincter and the levator ani muscles, which are innervated by the pudendal nerve (the inferior rectal and perineal nerves), correspond to the abaxial rectus muscle. A recent study revealed that muscular branches diverged from the pudendal nerve innervate classical superficial and deep portions of the external anal sphincter, and that the subcutaneous part predominantly receives independent branches from the sacral plexus (Plochocki et al., 2016). Thus, our dual innervation model may explain the biological significance of the dual innervation patterns in the pelvic floor.

Compared to that the levator ani and external anal sphincter muscle come to the table of discussion for such dual innervation, the perineal muscles, which belong to the abaxial compartment (Valasek et al., 2005), are exceptions and receive the solo innervation by the branches of the pudendal nerve. In our model, the pudendal nerve is the homolog to the canonical intercostal nerve, and is responsible for the innervation of the abaxial muscles associated with the perineal membrane (Figure 6). Therefore, our model also explains the exclusive innervation to the perineal muscles by the pudendal nerve.

General descriptions about the terminal branches of the pudendal nerve are that they bifurcate into the deep branch, the dorsal nerve of the phallus and superficial branch, and the superficial branch, the perineal nerve (Standring, 2015; Plochocki et al., 2016). The dorsal nerve of the phallus runs deep and anteriorly through the perineal membrane to the glans. The perineal nerve further travels anteriorly, passing between the superficial and deep transverse perineal muscles, runs superficially to the surface of the perineal membrane, and terminates as the posterior scrotal (labial) nerve, following the innervation to the perineal muscles. Because of the analogy that the intercostal nerve runs between the inner and innermost intercostal nerves, we allocated the superficial transverse perineal and cavernous perineal muscles to the internal oblique muscle class, and the deep transverse perineal muscle to the transverse muscle class (Figure 6, Table 1). The perineal nerve, which runs along the terminal end of the pudendal artery, mirrors the intercostal nerve that runs along the intercostal artery. This supports our hypothetical schema that the deep branch of the perineal nerve is the homolog of the intercostal nerve, and is responsible for the innervation to the perineal muscles.

Next, we allocated the ventral outer body wall cutaneous branch to the dorsal nerve of the phallus (terminal extension of the deep perineal nerve), based on the deep trajectory relative to the perineal membrane (Figure 6). The dorsal outer body wall cutaneous branch in the pelvic region corresponds to the inferior cluneal nerve. Finally, the posterior scrotum or labial nerve corresponds to the pelvic homolog of the ventral division of the lateral cutaneous branch of the spinal nerve in our model. The posterior femoral cutaneous nerve may partly contain the dorsal division of the lateral cutaneous branch. The above descriptions of the distributions of the three components are hypothetical, based on our model. Nevertheless, we are confident the above classification of nerve branches helps conceptualize the tangled networks of nerve branches in the pelvic floor.

#### 3 - 7) Muscular organization of the perineum

Hall et al. (2017) previously documented a homology of the muscular organization in the body wall between the pelvic and abdominal regions. We provide a different view of muscle homology between the pelvic and thoracic regions, based on the innervation patterns of the different classes of nerve branches. The homologous muscles between the two segments of the body wall are the following: the innermost intercostal muscle (transverse class, abaxial) to the deep transverse perineal muscle, the internal intercostal muscle (internal oblique class, abaxial) to the superficial transverse and perineal muscles, the external intercostal muscle (external oblique class, primaxial) to the coccygeus muscle, and the subcostal muscle (longus transverse class, primaxial) in the thoracic cage to the obturator internus. The subdivisions of the external anal sphincter and levator ani muscles correspond to the respective rectus abdominis muscle in the primaxial and abaxial compartments, according to the innervation by the pudendal nerve or direct branch from the sacral plexus.

The body wall organization of the perineum is different from that of other regions. The anterior portion of the perineum has an additional body wall, the perineal membrane, to the wall of the levator ani muscle (Standring, 2015). In our model, all muscles associated with the perineal membrane belong to the abaxial compartment and receive preferential innervation of the pudendal nerve, which contains the outer body wall and the anterior abaxial branch (homolog to the intercostal nerve). On the other hand, muscles around the anal canal belong either to the primaxial or abaxial compartments, and thus receive both innervations of the direct branches and pudendal nerve. Given the continuation between the external anal sphincter and perineal muscles, as described in the previous chapter, the innervation pattern in the pelvic floor suggests that the perineal membrane may be the anterior expansion of the abaxial compartment, reminiscent of the lower abdominal region. However, the difference is that the direct muscular branches exist in the perineum, but not in the lower abdominal region. The pudendal nerve may release the primaxial element of nerve branches to the rectus muscle derivatives (anterior primaxial branch) at the proximal segment because the pudendal nerve is exclusively destined to the abaxial compartment (perineal membrane). Thus, we attributed the separation of the primaxial anterior branch from the pudendal nerve to the unique body wall organization of the anterior perineum and possibly to the formation of the perineal membrane as a result of cloacal septation.

#### 3 - 8) Cervical region

The muscular organization of the neck is significantly different from that of the other body regions; the neck mostly lacks wall muscles, such as oblique and transverse muscles. This makes it challenging to allocate the named nerve branches to the three nerve components in our model, because there are no landmark intermuscular spaces.

However, there are rectus muscle derivatives in the neck, which are also in a two-layered structure, as in the pelvic region. The superficial long rectus muscles bridge over the thyroid cartilage from the sternum or scapula to the hyoid bone (sternohyoid and omohyoid), and the deep short rectus muscles connect the thyroid cartilage to the hyoid bone (thyrohyoid) or the sternum (sternothyroid). We allocated the surface rectus muscles (sternohyoid and omohyoideus) to the primaxial ones (superficial rectus class) and deep muscles (thyrohyoid and sternothyroid) to the abaxial ones (deep rectus class; Table 1). Because the diaphragm, is considered as a deep rectus muscle, the phrenic nerve is equivalent to the canonical intercostal nerve homolog (Table 1).

In the cervical region, from the C3 to C8, the scalene muscles (anterior muscle in the ventral subgroup, and medial and posterior muscles in the dorsal subgroup), longus capitis, and longus colli are innervated by branches (non-plexus forming branches) from the root of the spinal cord, and thus comprise the primaxial muscle group in our model (Figure 7). The deep prevertebral fascia enwraps these muscles around the vertebrae together with the intrinsic back muscles, and may therefore demarcate the primaxial compartment in the cervical region (Linder, 1986; Guidera et al., 20143; Kitamura 2018).

**Figure 7.**
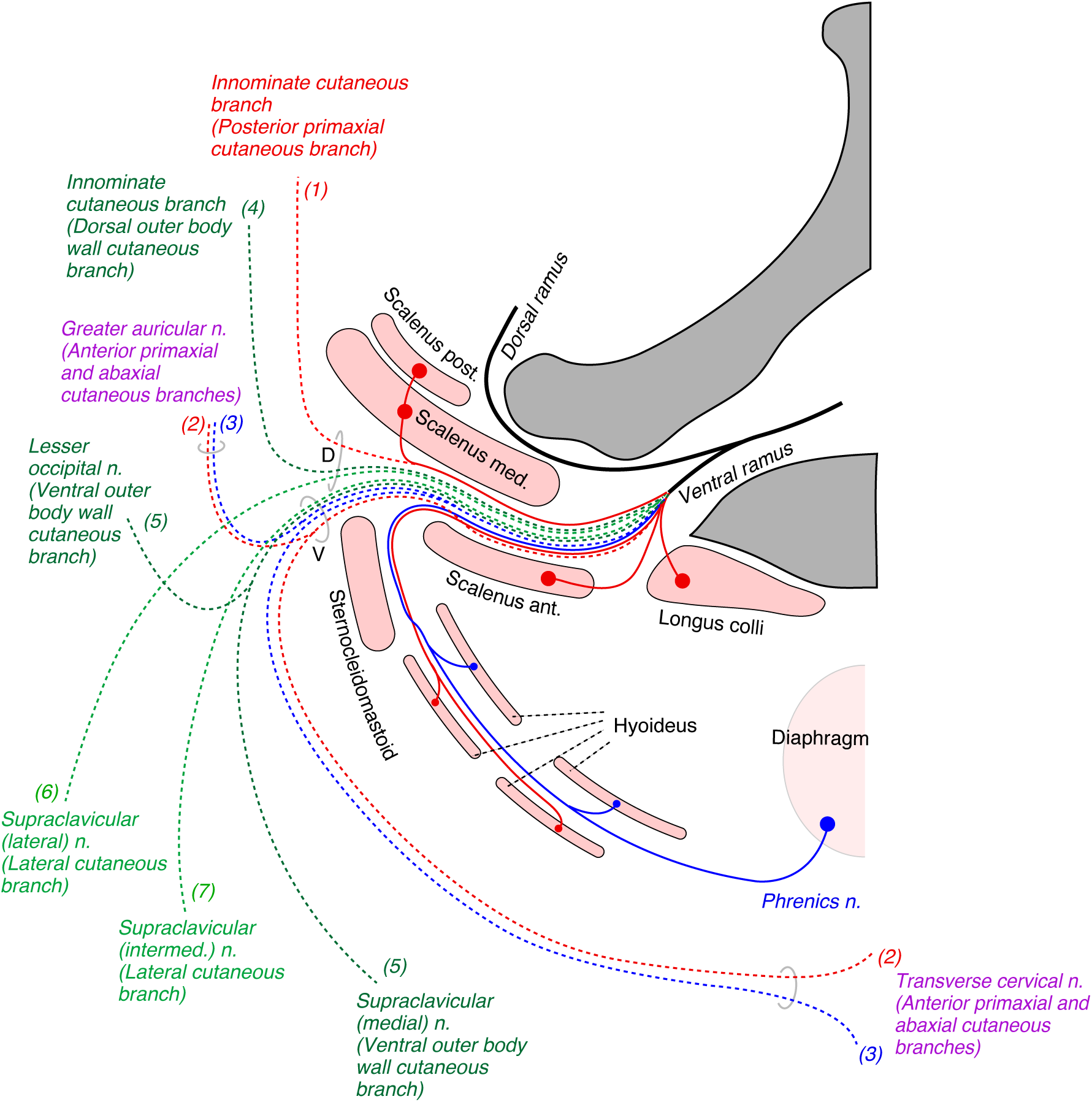
Hypothetical distribution of primaxial and abaxial-responsible nerve branches in the neck region. The positions of all branches and muscles are projected to a mid-transverse plane of the neck. The primaxial motor branches (red lines) include direct twigs to the scalene, longus colli, and superficial infrahyoid muscles. The abaxial motor branches (blue lines) innervate deep infrahyoid muscles and the diaphragm. The seven predicted cutaneous branches are numbered accordingly to our model. The posterior primaxial cutaneous branch (1) is innominate. Two anterior branches or the primaxial (2, red dashed lines) and abaxial compartments (3, blue dashed lines) are in the same bundle, and are bisected into the greater auricular and transverse cervical nerves. The dorsal outer body wall cutaneous branches (4) are innominate. The ventral outer body wall cutaneous branch (5) splits into the lesser occipital and medial branch of the supraclavicular nerve. The dorsal (6) and ventral divisions (7) of the lateral cutaneous branch are the lateral and intermediate branches of the supraclavicular nerve, respectively.

With regards to sensory innervation in the cervical region, the three predicted classes of sensory branches cannot be accurately assigned to the known cutaneous nerves. As stated previously, because there are no muscles encasing the neck wall, all cutaneous branches run without the reference to surrounding muscles. Furthermore, the sensory branches from the upper cervical spinal cord are split into ventrally directed (the transverse cervical and supraclavicular nerves) and dorsally directed (the lesser occipital and greater auricular nerves) groups; nevertheless, all the branches sprout from the ventral aspect of the spinal roots. This bisection makes allocation more difficult.

For the reasons above, we postulate that the cervical cutaneous nerves are composite. The transverse cervical nerve, lesser occipital, and greater auricular nerves often duplicate or even triplicate (Gupta et al., 2013; Ravindra et al., 2014). This numerosity suggests that the above cutaneous nerves may contain diverse nerve elements that may not be necessarily fasciculated under certain conditions. We also postulate undescribed cutaneous branches in the posterior cervical triangle of the neck. Some of these nerve branches are already known in other primates (Kato et al., 1990). With these postulations, we feel that it is adequate to apply our model to the gross anatomy of the cutaneous nerve in the neck region, based on the ramification and distribution patterns (Figure 7).

The transverse cervical and greater auricular nerves are ventrally and dorsally oriented pairs of anterior primaxial and abaxial branches (Figure 7). The lesser occipital and medial supraclavicular nerves are the ventral component of the ventral outer body wall cutaneous branches. The lateral and intermediate supraclavicular nerves are the dorsal and ventral components of the lateral cutaneous nerve. We propose the existence of the dorsal components of the outer body wall and posterior primaxial anterior cutaneous branches further dorsal to the lateral supraclavicular nerve (innominate branches in Figure 7).

#### 3 - 9) Brachial region

The innervation targets of the spinal nerves from the C5 to Th1 axial levels are all devoted to the upper appendicle (Figure 8). The mainframe of the brachial plexus corresponds to the outer body wall nerves, which innervate most of the upper appendicular muscles. The outer body wall branches, such as the pectoral and dorsal thoracic nerves, from the brachial plexus innervate the pectoral muscles (ventral group) and the latissimus dorsi (dorsal group), respectively. We described the sensory termination of the ventral outer body wall branch in the previous chapter.

**Figure 8.**
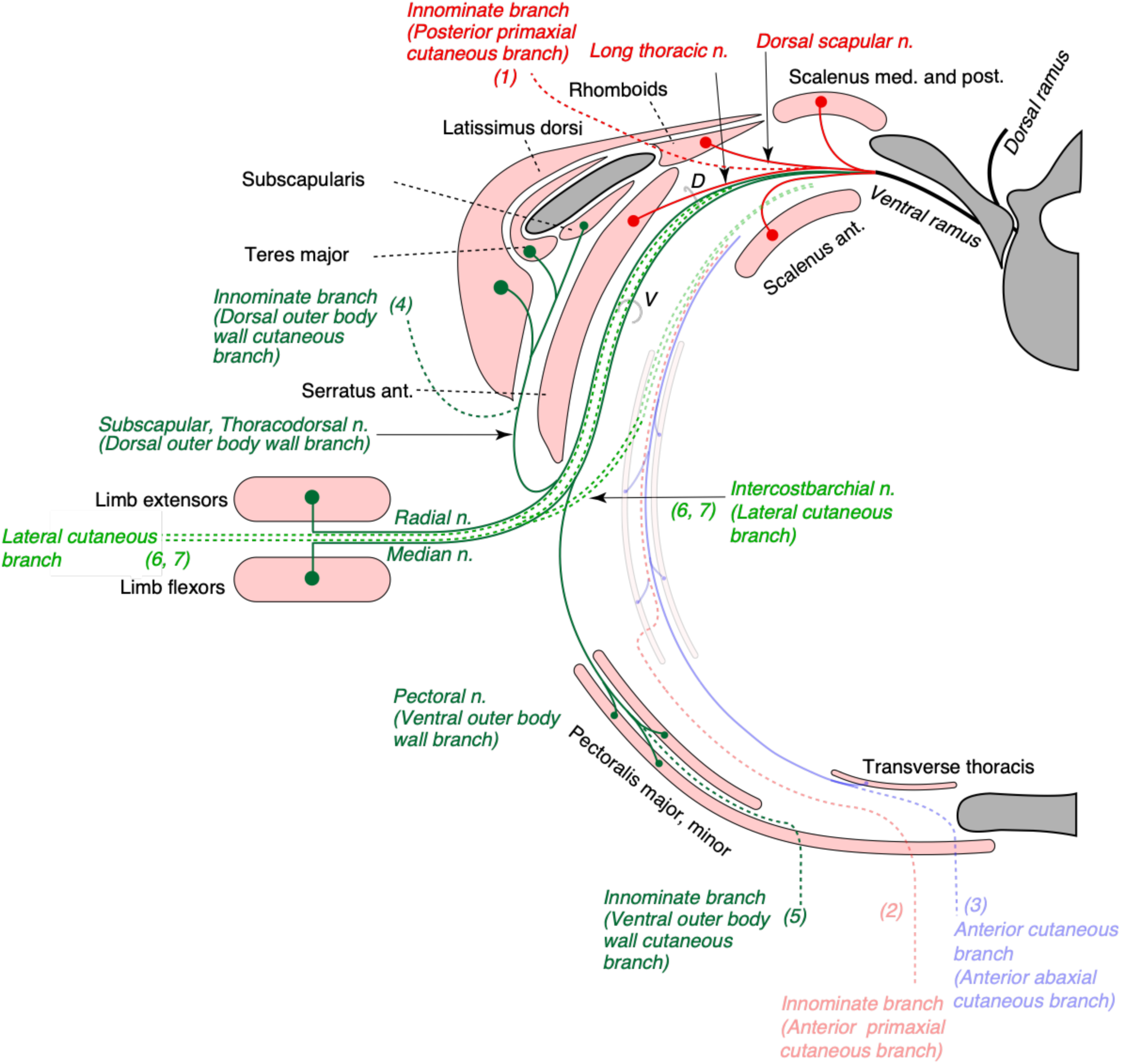
Hypothetical innervation pattern at the brachial level. Nerve branches are projected onto a transverse plane around the level of the fifth thoracic vertebra. Because the branches from the brachial plexus do not innervate muscles within the thoracic body wall, the anterior cutaneous branches and muscles in the thoracic wall are in opaque colors. The only exception to this is the intercostobrachial nerve from the Th1 (light green, 6, 7). The primaxial compartment-responsible nerves (red lines) are direct branches to the scalenus muscles, the long thoracic nerve to the serratus anterior muscle, and the dorsal scapular nerve to the rhomboids. The posterior primaxial cutaneous branch (1, red hatched line) is innominate. The outer body wall branches are dark green lines, and innervate the latissimus dorsi (thoracodorsal nerve), subscapularis (subscapular nerve), teres major (thoracodorsal nerve), and pectoral muscles (pectoral nerves). The lateral cutaneous branch (6 and 7) carries motor innervations, such as the radial and median nerves (outer body wall nerve, dark green), into the upper limb. The cutaneous extensions of the outer body wall branches (4, 5) are innominate.

The levator scapular, rhomboid, and anterior serratus muscles, which belong to the muscle class of the external oblique in the primaxial compartment, receive innervation by the dorsal scapula, or long thoracic nerves, which diverge from the point just proximal to the cord segment of the brachial plexus (Figure 8). The segmental branches to these three muscles do not maintain individuality and take shape as combined nerves. The union of the branches highlights differences in these two nerves from the other segmental branches diverged from the proximal root segment. True segmental branches innervate the scalene muscle (anterior scalene in the longus-transversus class, and middle and posterior scalene in the external oblique class).

Concerning sensory branches from the brachial plexus, the nerve branches piercing the rhomboid muscle from the ventral ramus distribute the lateral portion of the back (Aizawa and Kumaki, 1996). These reported nerves may correspond to the posterior primaxial and dorsal outer body wall cutaneous branches (Figure 8). On the other hand, because no muscle or dermis is associated with the ventral body wall of the brachial region in our model, there are no branches equivalent to the canonical intercostal nerve, anterior primaxial, or ventral outer body wall branches (Figure 8).

### 4. Validation of our model by experimental observations in the embryonic mouse

#### 4 - 1) Analysis of embryonic branching pattern in the body wall

Our model predicts that the single intercostal nerve trifurcates into superficial, deep, and lateral cutaneous branches, the first two of which reach the anterior midline of the thoracic region during early developmental periods. We tested this prediction using whole-mount transparent specimens of the thoracic walls from mouse embryos, in which we immunohistochemically visualized all neuronal branches.

In the thoracic walls of mouse embryos at embryonic day (E) 13, the intercostal nerves trifurcated into the lateral cutaneous, superficial, and deep branches at some length from the root (Figure 9). The lateral cutaneous branch always took the most superficial position within the thoracic wall, and sprouted anteriorly directed branches in addition to the proper branch to the dermis. The spinal nerve from Th1 always lacked the lateral cutaneous and deep branches. In the spinal nerves from the upper thoracic levels, especially in Th2 and 3, the superficial branch was often absent.

**Figure 9.**
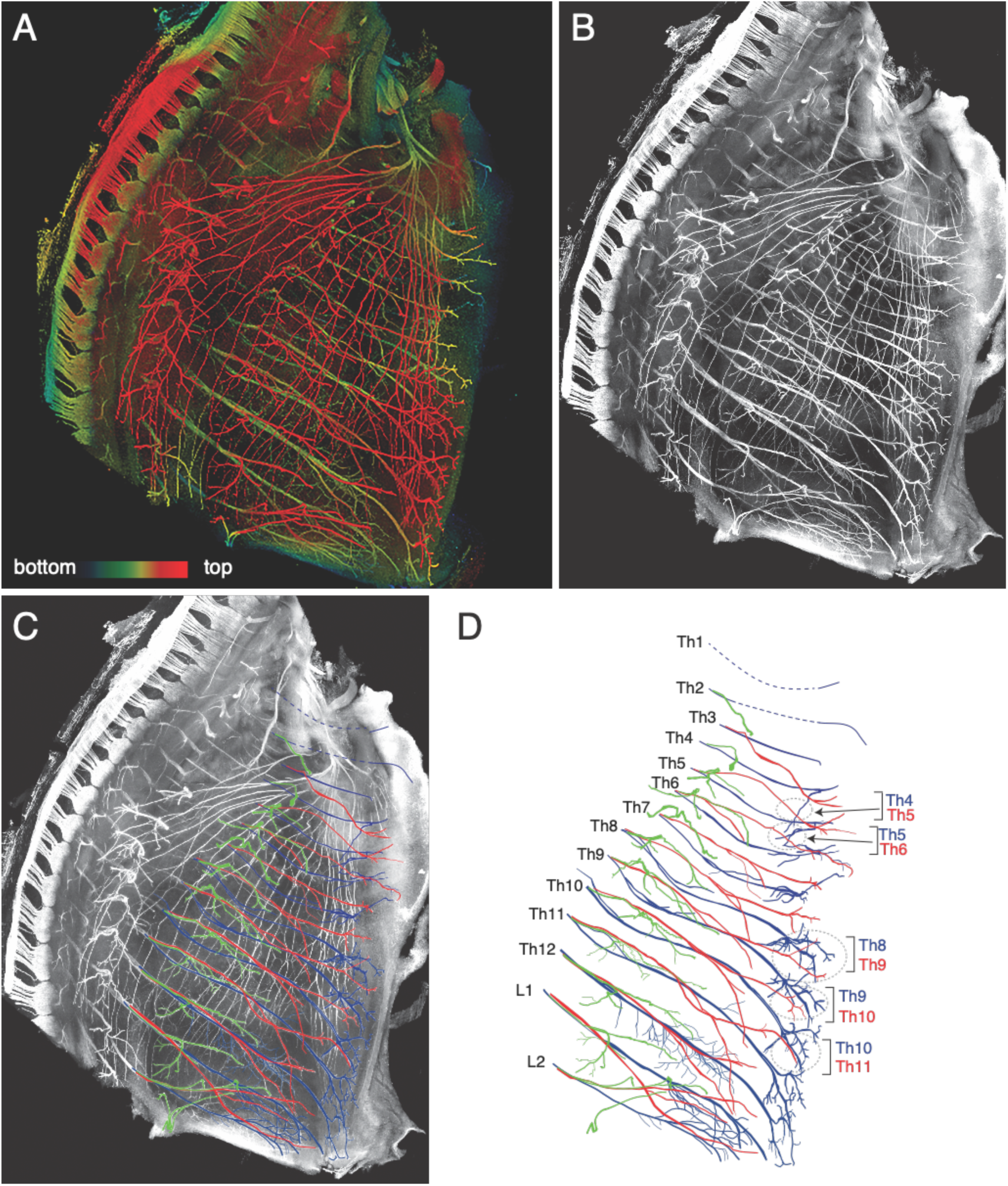
Branching pattern of the intercostal nerve in a transparent thoracic wall from a mouse embryo on E13. All micrographs are the unilateral side of the thoracic wall, and are in surface view. The dorsal side is toward the left of the micrographs. The anterior side is toward the top of the micrographs. (A) MIP image. Nerve branches are color-coded; those close to the outer surface are colored in red, and those close to the parietal surface are in blue. (B) A micrograph of a black and white MIP image without depth information. The vertically running branches are for the dermal panniculus muscle from the brachial plexus (Petruska et al., 2014). (C) Traced intercostal nerves; the deep branches (blue), superficial branches (red), and lateral cutaneous branches (green). When the surface skin was peeled off, the terminal branches of the lateral cutaneous branches were also removed. No tracing of the branches for the dermal muscle is included. (D) Illustration of the traced branches. Blue, deep branches; Red, superficial branches; and Green, lateral cutaneous branches. The hatched circles indicate overlapping innervations by the nearby superficial and deep branches.

The superficial branch from an axial level and the deep branch from the one above the axial level often innervated the same areas around the ventral midline, albeit the overlapping did not necessarily exist at all axial levels (Figure 9D). The overlapping innervation by spinal nerves from the two nearby axial levels indicates that the superficial and deep branches may receive axial specifications in different manners. Hence, one branch is not merely the collateral branch from the other branch; rather, they are likely to be distinctive from each other.

The branching pattern in slightly older embryos (E15) was different from that on E13, in that the superficial branch became significantly thinner in diameter and shorter in length, and does not reach the ventral midline (Figures 10 and 11). Instead, the anterior branches that sprouted from the lateral cutaneous branch were notably conspicuous, especially in the lower thoracic levels. Programmed cell death occurs among spinal motoneurons of mouse embryos from E13 to E15 (Lance-Jones, 1982; Yamamoto and Henderson, 1999). The time-coincident events between the neuronal origins and processes indicate that the size reduction of the superficial branch may result from the death of the parental motoneurons of the superficial branches.

**Figure 10.**
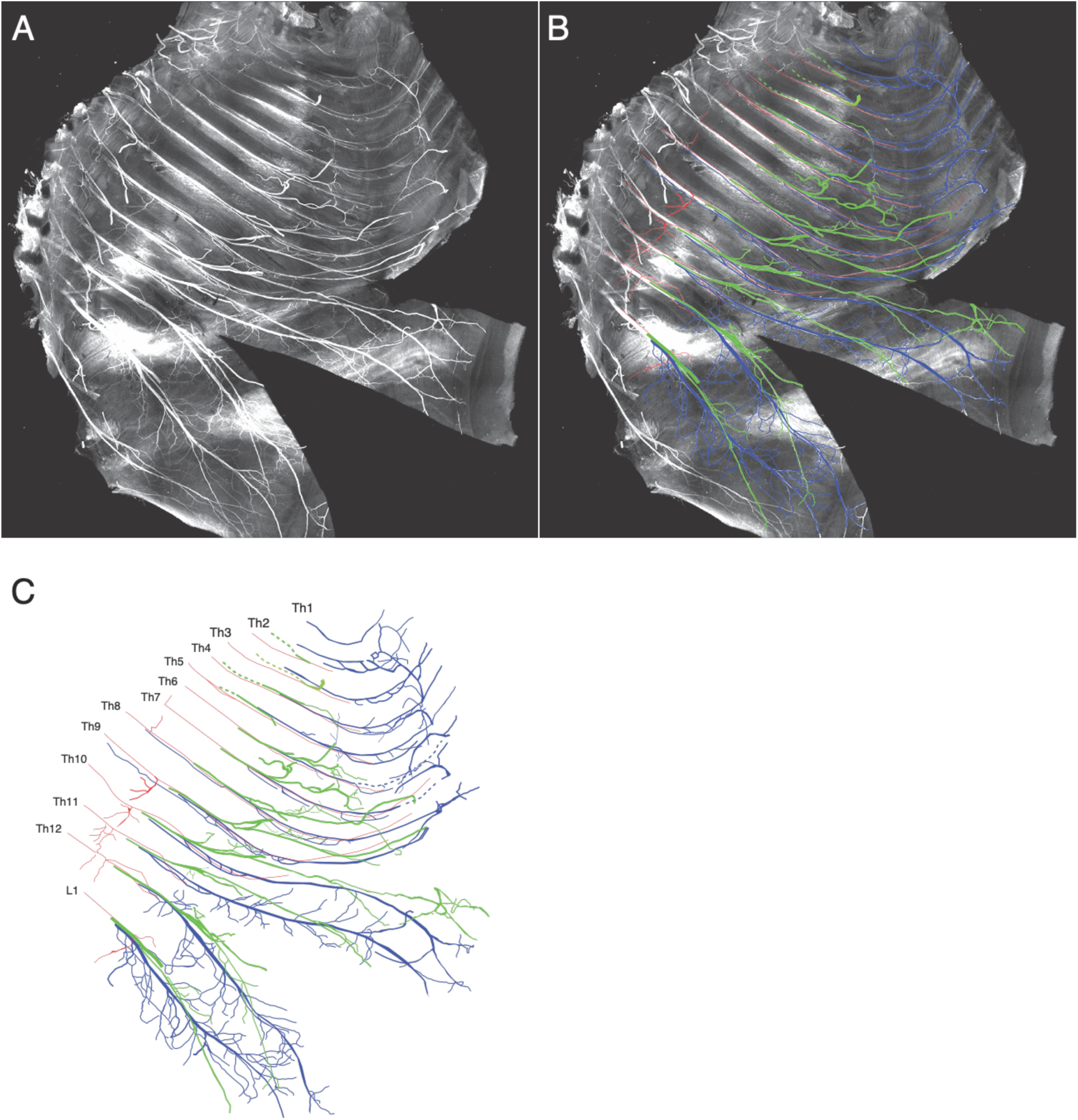
Branching pattern of the intercostal nerve from a mouse embryo on E15. All micrographs are in the surface view of the unilateral side of the thoracic wall from an E15 embryo. The dorsal side is toward the left of the micrographs. The anterior side is toward the top of the micrographs. Because of the conspicuous curvature of the thoracic wall, two split lines are in the lower portion of the wall for mounting. (A) MIP image. (B and C) Tracing of the branches. The color code: blue, deep branch; red, superficial branch; and green, lateral cutaneous branch.

**Figure 11.**
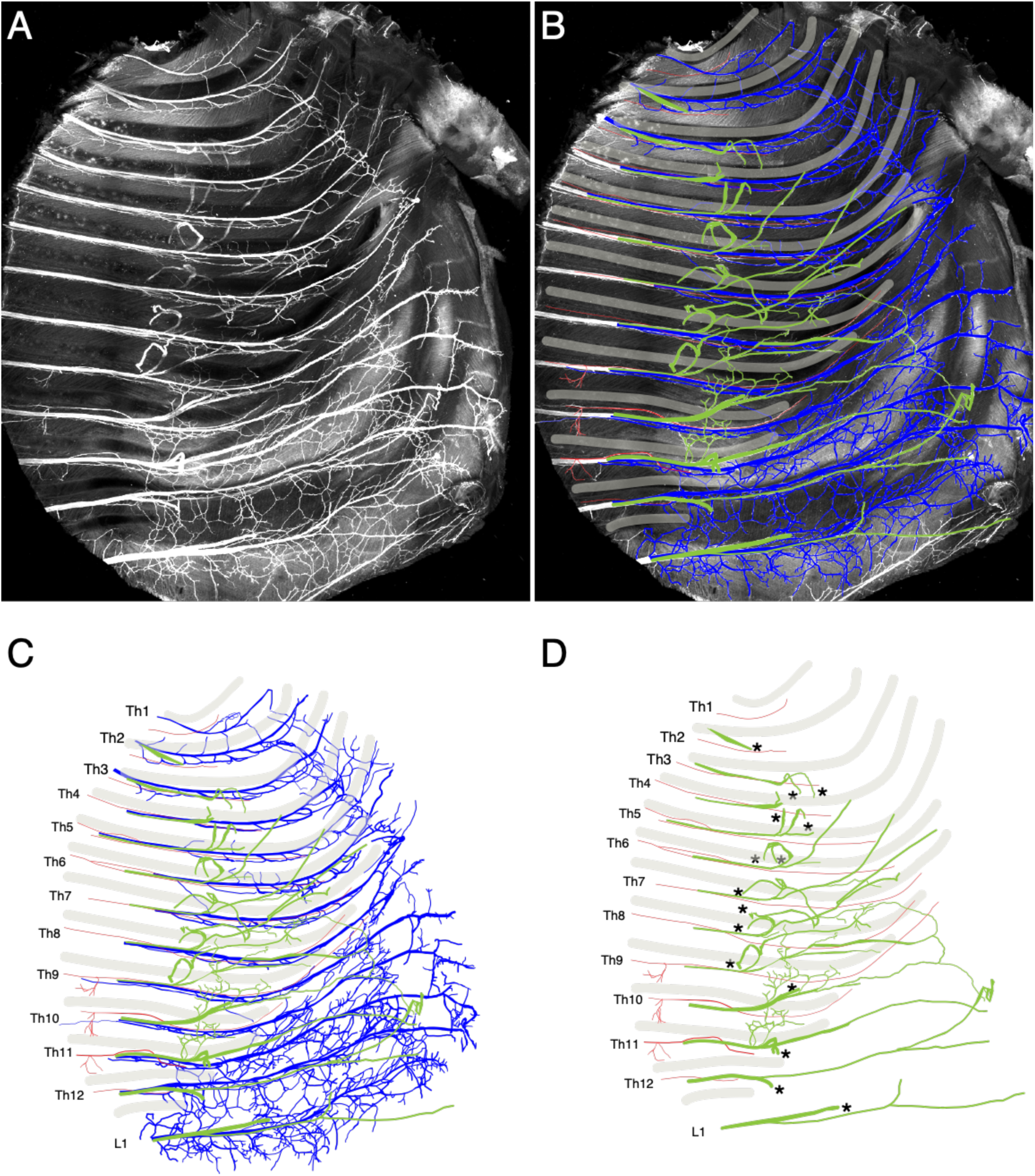
Close-up view of the thoracic wall from an E15 embryo. The anterior side is toward the top of the micrographs. The costal side is toward the right of the micrographs. The cartilaginous costae are indicated by gray bars. (A) Black and white MIP image. (B and C) The superficial branches (red) run along the lower margin of the costae while the deep (blue) and lateral cutaneous (green) branches change their courses toward one or more of the above intercostal spaces. (D) The superficial and lateral cutaneous branches are extracted for comparison, showing the segmental and non-meristic innervation patterns of the superficial and lateral cutaneous branches, respectively. Asterisks are stumps of the dermal branches of the lateral cutaneous branch, because the skin and dermis were removed for transparency.

The deep branches traveled along the lower margins of the costae toward the anterior midline, where their terminal branches that originated from different axial levels communicated with each other (Figure 11C). Similar communication of the thoracolumbar nerve in the human abdominal wall has also been reported (Rozen et al., 2008). The lateral cutaneous branch proper (dermal branch) and the anterior branches of the lateral cutaneous branches changed their running-directions around the middle of the costae (Figures 11C and D). They pointed toward one intercostal space above or even more to the anterior thoracic region, and often united with the nearby anterior branches. However, the shortened superficial branches were meristic, and remained within the intercostal space at the same axial level (Figures 11C and D). Thus, the superficial branches maintain their axial constraint, whereas the deep and lateral cutaneous branches are free from the axial confinement. Because the axial specification by Hox-code differs between the somite and the lateral plate, there must be a mechanism by which coordination between them is accomplished for global body wall patterning (Cohn et al., 1997; Nowicki and Burke, 2000). Nerve branches from the ventral rami are a structure that requires a specific coordinating mechanism because the peripheral nerve serves as a connection between the primaxial and abaxial compartments. The trajectories free from the original axial specifications of the deep and lateral cutaneous branches indicate that they may be responsible for the innervation of the abaxial compartment by bridging two different axial environments. On the other hand, the superficial branches remain within the same intercostal space as with the original axial specification, and may innervate the structures in the primaxial compartment by complying with its axial restriction. We also would like to point out that the meristic nature of the superficial branch fits our working hypothesis that the nerve branch responsible for the abaxial compartment is segmental.

The above-observed innervation patterns in the embryonic periods are entirely in favor of our model. Nakao and Ishizawa (1994) reported a similar branching pattern of the intercostal nerve, in which the ventral rami of the spinal nerves diverged into the three primary branches, including the branch to the external intercostal muscle (superficial intercostal nerve), lateral cutaneous branch, and anterior branch of the intercostal nerve. The differences between their study and ours are that they did not confirm that the superficial branch reached the anterior midline from E13 embryos, and also that the anterior branches sprouted from the lateral cutaneous branch proper. Because they used the histochemical method to visualize the motor nerves, it was challenging to demonstrate the entire innervation patterns of the intercostal nerve in their study. Because no study on the development of the intercostal nerve has been carried out since the rise of current tissue-clearing methods, we do not have information on the embryonic intercostal nerve based on a clear image of the entire branching pattern with special attention for branches in the deep portion.

However, even after the development of tissue-clearing methods, few studies have investigated the development of the intercostal nerve. The mouse embryos on the front cover of Developmental Biology by Gilbert and Barresi (2016) and Figure 1A in the paper entitled “Frizzled3 controls axonal development in distinct populations of cranial and spinal motoneurons” by Hua et al. (2013), exhibited the same branching pattern as that seen in our observations, which are three radiated branches from the ventral rami. Many who are interested in the spinal nerve may interpret either one of the branches as a collateral of the intercostal nerve. However, based on the detailed information on human gross anatomy and the view of the lateral somitic frontier, we adequately interpreted both branches as principal components of the intercostal nerve.

#### 4 - 2) Analysis for the expression of a motoneuron marker following backtracing

##### 4 - 2 - 1) Injection into putative primaxial muscles

Our working hypothesis that the nerve branches to the primaxial muscles from the ventral rami share the same traits as branches of the dorsal rami. We also tested this hypothesis at a microscopic level, regarding marker expressions in the parental motoneurons of segmental branches from the ventral rami. The motoneurons that innervate the back muscles exist in the medial portion of the medial motor column (MMCm), and express a unique marker of the LIM-class homeobox gene, Lhx3 (Tsuchida et al., 1994, Dasen et al., 2008; Rousso et al., 2008). Accordingly, we analyzed the Lhx3 expression pattern in the motoneurons that innervate putative primaxial muscles in our model using the retrograde labeling method. Dextran-Alexa594 was injected into the longus colli, anterior scalene, anterior serratus, psoas, and quadratus lumborum. All labeled motoneurons innervating these muscles (except for the anterior serratus) expressed Lhx3 in the medial side of the medial motor column (MMCm) in E13 or E15 mouse embryos (Figures 12A–E). The expression levels of Lhx3 in the labeled motoneurons are often lower, compared to surrounding Lhx-3-positive cells. The labeled motoneurons to the anterior serratus muscle also expressed Lhx3, but in the most lateral portion of the ventral horn (Figure 12C). In a previous study, the motoneurons innervating the rhomboid muscles are also in the most lateral side of the ventral horn (Tsuchida et al., 1994). The distinctive location of the motoneurons may reflect the unique nature of the nerve branches to the rhomboid and anterior serratus muscles. Segmental branches from different axial levels to these muscles do not maintain the meristic nature throughout their course, and unite to form the dorsal scapular and long thoracic nerves. Due to technical reasons, we could not analyze the motoneurons to all muscles selected as primaxial muscles in our model. Nevertheless, the above Lhx3 expression pattern is also in favor of our model at a molecular level.

**Figure 12.**
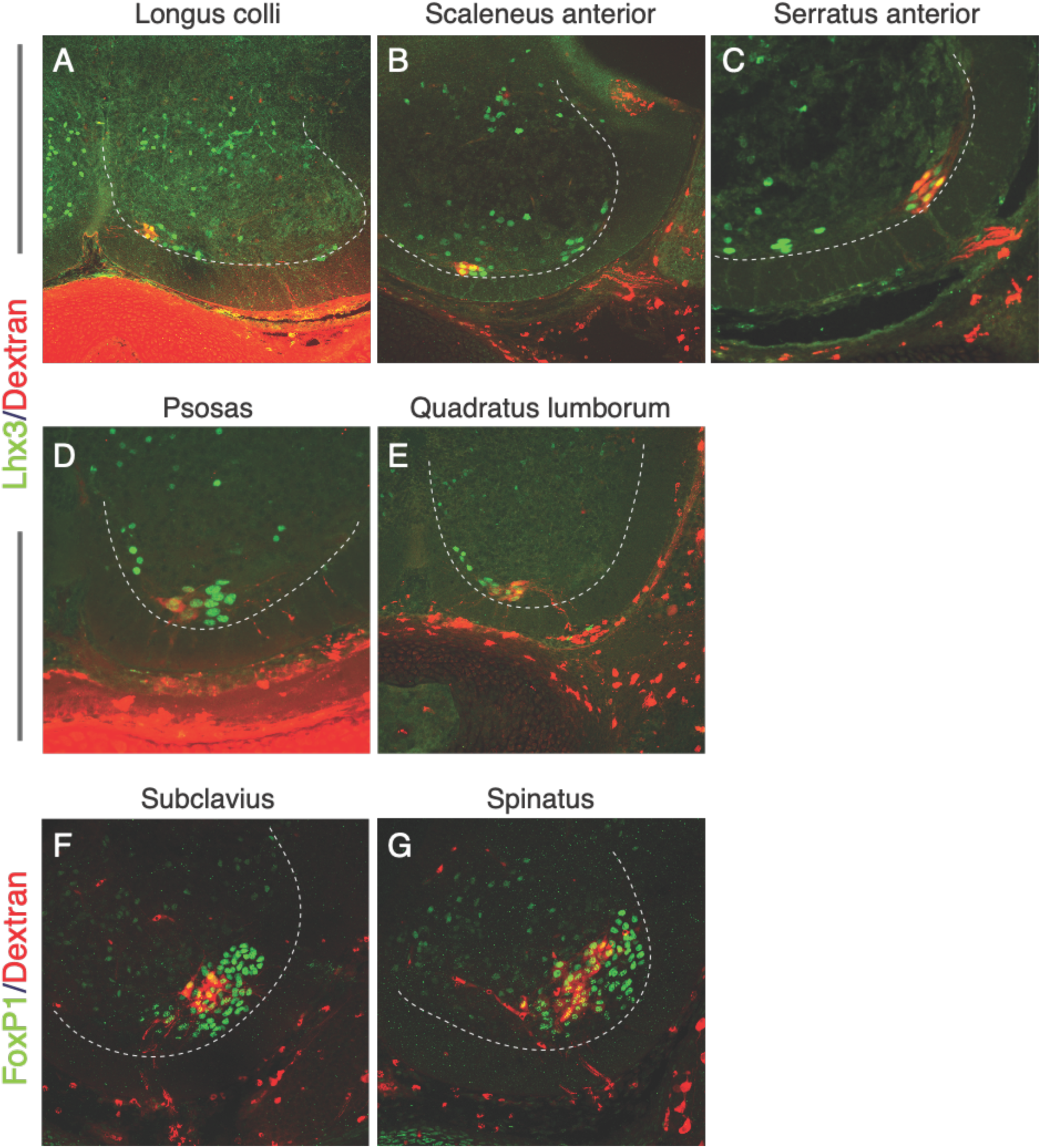
Marker expression patterns after the retrograde labeling. Lhx3 expression (A to E: green nuclei) and FoxP1 (F and G: green nuclei) in labeled motoneurons (red) following the injection of the tracer into the target muscles. The ventral quadrant of the brachial and lumbar spinal cord from E13 or E15 mouse embryos are shown, and dashed lines mark the ventral horns. In all micrographs, the midline and the dorsal side are toward the left and top, respectively. (A) Longus colli (E13). (B) Scalenus anterior (E13). (C) Serratus anterior (E13). (D) Psoas (E13). (E) Quadratus lumborum (E13). (F) Suvclavius (E13). (G) Spinatus (E13).

The primaxial muscles, such as the longus colli, connect the cervical vertebrae and act on the body axis. Because of these positional and functional similarities to the intrinsic back muscles, the parental motoneurons may express a medial motor column (MMCm) marker, Lhx3. Similarly, the Lhx3 expression in the motoneurons innervating girdle muscles, such as the serratus anterior muscle, may come from the close location of the target muscles to the body axis. We also tested this possibility of whether Lhx3 is a marker for motoneurons innervating the girdle muscles. The tracer injection into the supinatus and subclavius muscles revealed that the motoneurons innervating these muscles were FoxP1-positive, a marker for LMC motoneurons (Dasen et al., 2008; Figures 12F and D).

##### 4 - 2 - 2) Injection into the superficial and deep branches of the intercostal nerves

We next injected the tracer into the superficial and deep branches of the lowest four intercostal nerves in E13 embryos. The injection was made at the site as close as possible to the putative location of the rectus muscle, although the rectus muscle itself was almost discernible (Figure 13A). After the injection into the superficial branches, we found that the labeled motoneurons were in the medial side of the MMC (MMCm) and Lhx-3-positive (Figure 13B). Following the injection into the deep branches, the labeled motoneurons were in the lateral side of the medial motor column (MMCl), and some of the labeled motoneurons expressed Oct6 (SCIP), which is known as a marker for phrenic motoneurons (Figure 13C; Rousso et al., 2008). However, most of the labeled motoneurons were Oct6-negative. We have never observed Lhx3-positive labeled motoneurons in the MMCm following the injection into the deep branches (Figure 13D).

**Figure 13.**
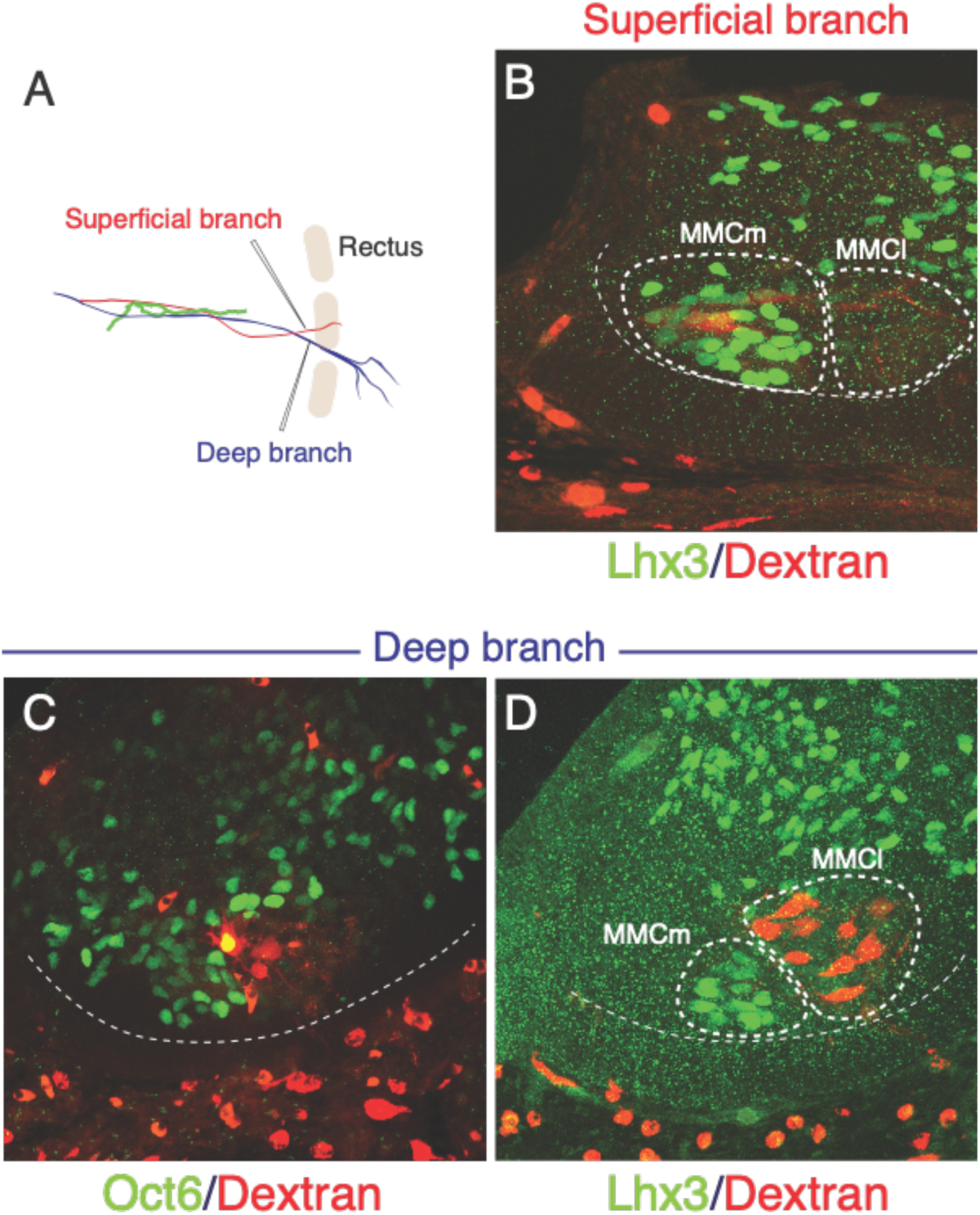
Differential origins of the superficial and deep branches of the intercostal nerves. All micrographs are transverse sections of the lower thoracic spinal cord from E13 embryos. The ventral margins of the ventral horns are in thin dashed lines. The midline of the spinal cord is toward the left side, and the dorsal side is toward the upside of the micrographs. Thick dashed lines encircle the medial (MMCm), and lateral (MMCl) halves of the medial motor columns. (A) Injection points are in the superficial (red) and deep (blue) branches, which are close to the rectus muscle band (light brown). (B) The labeling (red) is in Lhx3-positive (green nuclei) motoneurons, which are in the MMCm, but not in the MMCl following the injection into the superficial branches. (C and D) Following the injection into the deep branches, labeling is restricted within the MMCl, where Lhx3 expression is negative, and the Oct6-positive nuclei (green) with a high expression level is observed in the labeled motoneurons (red) in the MMCl.

The Lhx3 expression in the apparent non-paraxial muscle, the rectus muscle, indicates that we may expand the border of the target muscles for MMCm motoneurons to a broader category of muscles. Collectively taking the above results into consideration, we may obtain the adequate conception that Lhx3 is a marker for the primaxial muscle-innervating motoneurons.

##### 4-2-3) Prx1 expression in the rectus muscle

We also analyzed paired box-related 1 (Prx1), a marker for the connective tissue associated with the abaxial compartment, in the rectus muscle in E13 mouse embryos (Durland et al., 2008). The rectus muscle, which was identified by MyoD expression and morphology, was entirely Prx1-negative in the lower thoracic region (Figures 14A–D). However, in the lower abdominal region, the rectus muscle consists of Prx1-positive and negative portions (Figures 14E and F). The observed Prx1 expression indicates that the rectus muscle is not uniform regarding which compartment it belongs to, whether it be primaxial or abaxial. The thoracic rectus muscle belongs exclusively to the primaxial compartment, whereas the abdominal rectus muscle encompasses both compartments. This differential Prx1 expression in the rectus muscle supports the architecture of the rectus muscle in our model. Furthermore, based on our assumptions, we suggest that the Lhx3-positive labeled motoneurons in the MMCm following the injection into the superficial branch may correspond to those innervating the Prx1-positive rectus muscle (primaxial rectus muscle), and Oct6-positive motoneurons to the Prx1-positive rectus muscle (primaxial rectus muscle).

**Figure 14.**
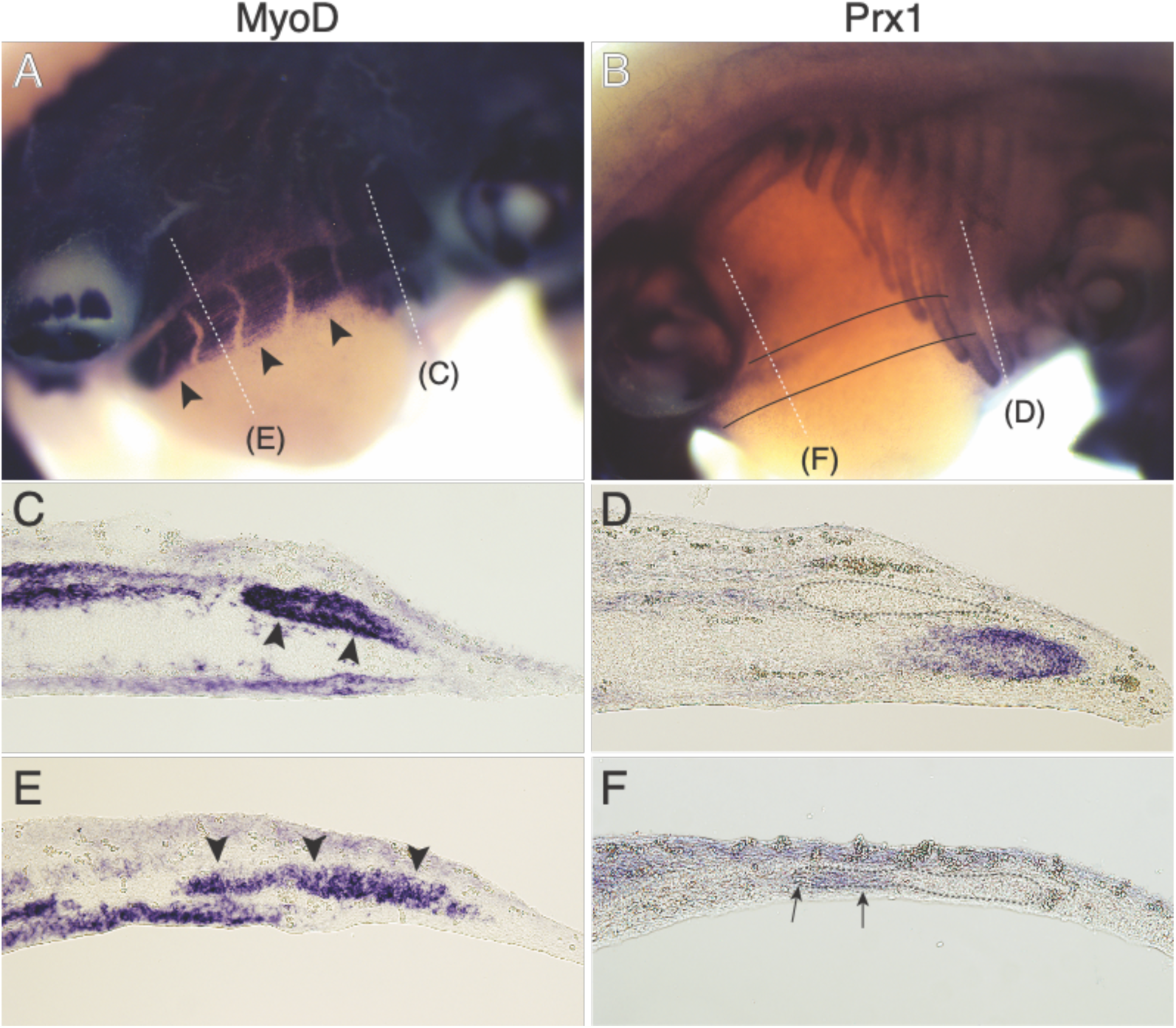
Expression patterns of MyoD and Prx1 in E13 mouse embryos. Micrographs in A and B are whole-mount in situ hybridization, and micrographs in C to F are sections cut from embryos processed for whole-mount in situ hybridization. White dashed lines in (A) and (B) indicate the approximate axial levels from which the sections originate. Arrowheads in (A), (C), and (E) indicate the rectus muscle. Black lines in (B) indicate the approximate position of the rectus muscle band. In (D) and (F), dashed lines encircle the locations of the rectus muscle, and arrows in (F) indicate Prx1 expression in the rectus muscle. The anterior midline and the surface of the body wall are toward the right and top of the micrographs of (C) to (F), respectively.

##### 4 - 2 - 4) Tracer injection into the infrahyoid muscles

The two-layered architecture of the rectus muscle is well preserved in the neck and pelvic regions, as described in the previous chapter. Because of easy accessibility to the embryonic neck region, we made the tracer injection into the infrahyoid muscles. However, because the individual muscles in the infrahyoid region were too small to be seen, the injection was made into the tissue mass below the hyoid cartilage and above the trachea. Our model predicts that motoneurons innervating the superficial infrahyoid muscles express Lhx3, and that those innervating the deep infrahyoid muscles express Oct6 (SCIP), which is now a marker for motoneurons innervating not only the diaphragm (rectus derivative) but also the putative deep rectus muscles. After the injection, we observed the labeling in the hypoglossal nucleus and the ventral horn. Within the ventral horn, some of the labeled motoneurons expressed Lhx3 or Oct6 (Figure 15). Lxh3-positive motorneurons were in the medial side of the ventral horn (MMCm; Figure 15A), and Oct6-positive motoneurons were relatively in the lateral side (Figure 15B). The Oct6-positive motoneurons were in the cluster of cells with high Oct6 expression (Figure 15B). This expression level indicates that Oct6-positive motoneurons are in the MMCl because the expression level is low in the MMCm (Rousso et al., 2008). As such, the expression patterns of Lhx3 and Oct6 in the cervical spinal cord also support our model. Rousso et al. (2008) reported that motoneurons with the MMCl profile (Oct6-positive) exist all along the axial levels. Our results, together with the Oct6 labeling in motoneurons following the deep branches of the intercostal nerve, indicate that Oct6 may be a marker for the motoneurons innervating the deep rectus class of muscle in the abaxial compartment.

**Figure 15.**
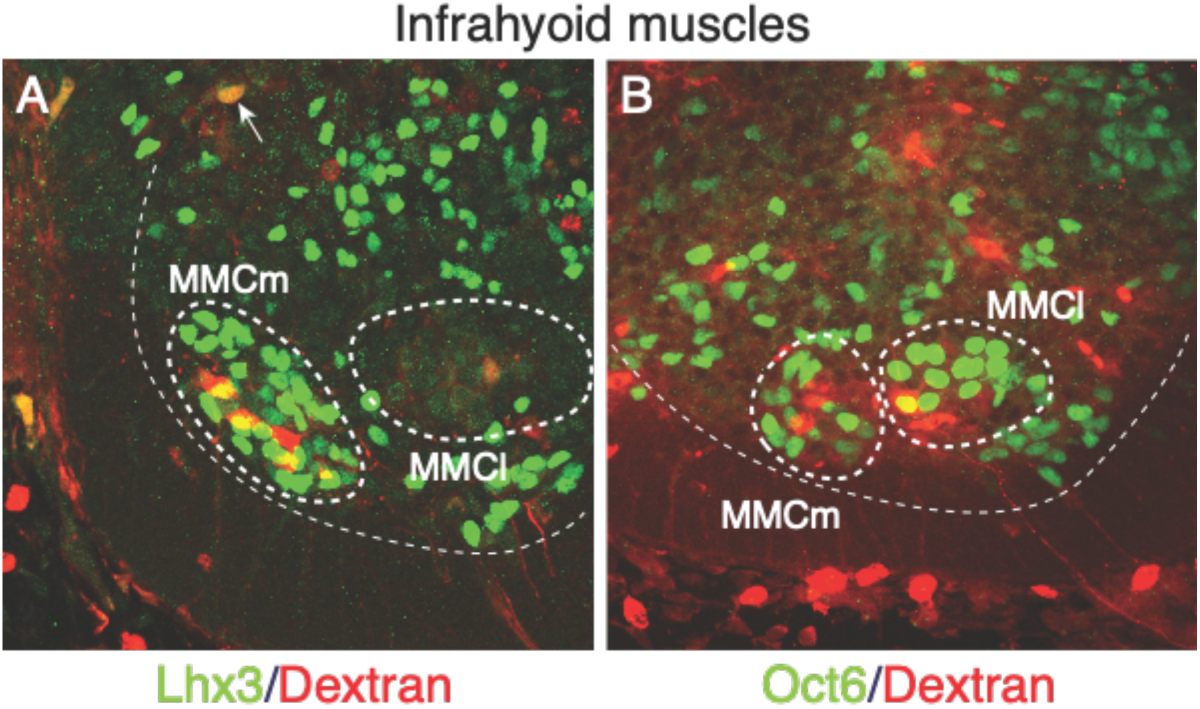
Lhx3 and Oct6 expressions following the injection into the infrahyoid muscles. All micrographs are transverse sections of the upper cervical spinal cord (C1 and 2) from E13 embryos. The boundaries of the ventral horns are thin dashed lines. Thick dashed lines encircle MMCm and MMCl. The midline of the spinal cord is toward the left side, and the dorsal side is toward the upside of the micrograph. (A and B) The labeled motoneurons are Lhx3-positive in the medial side of the medial column (MMCm) in (A) or Oct6-positive in the lateral side of the MMC (MMCl) in (B). The arrow in (A) indicates a labeled hypoglossal motoneuron.

##### 4 - 2 - 5) A proposed simple schema for the columnar organization of spinal motoneurons

Collectively, based on the above expressions of Lhx3 and Oct6 among the labeled motoneurons, we venture to put forward that more straightforward characterization of muscle varieties of innervation target in the three principal motor columns in the spinal cord. First, Lhx3-positive motoneurons, which are only in the MMCm, innervate muscles in the primaxial compartment. Lhx3-negative motoneurons, which are both in the MMCl and LMC, are for muscles in the abaxial compartment. The motoneurons in the LMC are FoxP1-positive and innervate the girdle and limb muscles. MMCl motoneurons are Lhx3- and FoxP1-double negative, and are for body wall muscles in the abaxial compartment. The MMCl motoneurons that innervate muscles only in the deep rectus class express Oct6 with high expression levels; thus, Oct6 is not a pan-MMCl marker. Accordingly, the three principal columns correspond to the neuronal origins of the three nerve components in our model; MMCm for primaxial compartment-responsible branches, MMCl for the homologous branch to the canonical intercostal branch responsible for abaxial body wall, and LMC for the outer body wall branch innervating the appendicular muscles.

Previously, Dasen (2009) proposed a plausible scenario of how three principal columns are evolved from the initial population of motoneurons. Firstly, MMCm is placed under the dominant actions of Lhx3; the appearance of a population escaped from the Lhx3 influence is preset to MMCl and LMC; finally, FoxP1 activity lays down the properties of LMC motoneurons on the evaded population. Our characterization of the motor column, based on the primaxial-abaxial distinction, is in accordance with the above scenario.

## CONCLUDING REMARKS

In this study, we proposed a model for the branching pattern of the spinal nerve, based on the view of the embryonic mesoderm compartments. Our model highlights the three-component composite of the spinal nerve, which differentially innervates the primaxial and abaxial parts of the body wall, as well as the appendicles. We believe that these findings could lead to radical changes in our current understanding of anatomical organizations of peripheral innervation patterns to the muscle and dermis, but consequently fills the gap between the real dissection and our perception derived from standard references. Primarily, explanatory notes on the branching pattern of the spinal nerve would fulfill the demands for information withstanding for practical use, especially in clinical medicine.

The embryonic branching pattern of the spinal nerve and marker expressions among spinal motoneurons support our model. Based on the embryonic observations, we further provided clues to a more straightforward view of the columnar organization of spinal motoneurons; the medial MMC for primaxial muscle-innervating motoneurons, the lateral MMC for motoneurons innervating abaxial body wall muscles, and the LMC for appendicular muscle-innervating motoneurons.

Also of significance is the evolutional aspect of the branching pattern. Since the primaxial and abaxial compartments are such fundamental patterning environments for the mesoderm derivatives, their selective expansion leads to the generation of evolutionary modifications in the vertebrate body plane. In our model, for example, the muscles controlled by primaxial-responsible branches constitute an essential set for respiration. Abaxial muscles in the body wall are also involved in active costal and abdominal breathing, but through more specific and efficient manners, including changing chest volume via the diaphragm, active exhalation by the inner intercostal muscle, and abdominal pumping by the oblique and transverse abdominis muscles. The selective innervation to the abaxial muscles by the respective branches is plausibly associated with the terrestrial adaptation, which requires the development of more sophisticated apparatus for breathing and locomotion. Thus, our model for the spinal nerve branching pattern would provide fundamental ground for comparative studies on evolutionary changes in the vertebrate body.

## MATERIALS and METHODS

### 1. Mouse embryos

We purchased pregnant mice from a local vendor and euthanized them under deep anesthesia with the isoflurane vapor. The uterus was dissected out and immediately soaked in ice-cold phosphate-buffered saline (PBS). Then, the mouse embryos were taken out in ice-cold PBS and stored in cold L-15 medium (Wako pure chemical) until use and immersed in cold 0.1M phosphate-buffered 4% paraformaldehyde (4%PA in 0.1M PB). The internal committee approved the protocol for the animal experiments in this study (Approval number 30071).

### 2. Retrograde-labeling

After the embryos were decapitated and eviscerated, target muscles and nerves were exposed in cold PBS, and Alexa594-conjugated Dextran (D22913, Invitrogen) was injected using a picosplitzer (IM300, Narishige). The injected embryos were kept in oxygen-bubbled DMEM (Wako pure chemical) for up to five hours at 30℃. After the incubation, the embryos were fixed in 4%PA in 0.1M PB for 5 hours and then immersed in 20% sucrose in PBS overnight. The appropriate segments of the spinal cord were cut out from the embryo and frozen-embedded in a mixture of two parts of 20% sucrose in PBS and one part of Tissue-Tek embedding media (Sakura, Japan).

### 3. Immunohistochemistry and image acquisition

Whole-mount neurofilament immunohistochemistry was carried out according to iDISCO protocol, except for the tissue-clearing methods (Renier et al., 2014). The primary neurofilament antibody was purchased from Biolegend (841001, 1:500 dilution).

For the simultaneous detection of the motoneuron markers in the labeled motoneurons, cryostat sections were cut and processed for immunohistochemistry as follows. The primary antibodies used in this study were Lhx3 (Abcam ab14555, 1:1000 dilution) and Oct6 (Abcam ab221964, 1:250 dilution). For Oct 6 immunohistochemistry, the sections were treated in 0.01M SSC buffer (pH = 7.0) for 20 minutes at 98℃ for antigen retrieval. The sections were blocked with 10% normal goat serum in PBS with 0.1% Tween 20 (PBST) for one hour, incubated with the primary antibodies overnight at 4℃, and then incubated with the Alexa498-conjugated anti-rabbit goat antibody (Invitrogen) at room temperature for 3 hours.

Images were obtained under a confocal microscope (A1, Nikon). For 3D reconstruction from captured images, Flurorender (Wan et al., 2009) and Adobe Photoshop applications were used, and maximum intensity projection images were created.

### 4. Tissue-clearing with TDE

The organic solvents used in the iDISCO protocol, such as tetrahydrofuran and dibenzyl ether, are toxic and require special care for handling. For this reason, we used 2,2′-Thtiodiethanol (TDE) for tissue clearing and mounting media (Staudt et al., 2007). After the immunohistochemistry, the samples were fixed in 4%PA in 0.1M PB for 30 minutes at room temperature. Following a brief wash with PBST, the samples were immersed in a graded series of TDE/water mixture with the final concentration of 98%, and mounted in a hand-made chamber. We made the chamber on a slide glass by stacking square vinyl frames, which were initially developed for in situ hybridization (AB-0578, Thermo Scientific). The depth of the room was adjusted by the number of stacking layers according to the thickness of the specimens. After placing the processed tissue samples, the chamber was filled with TDE and sealed with a coverslip. The specimen was therefore flattened and immobilized between the slide glass and coverslip.

Besides safety, the most significant reason why we used TDE, instead of dibenzyl ether, in the original iDISCO protocol, was to avoid refractive index mismatch along the light path. One of the leading causes to hamper the obtaining of bright and shaped images from deep inside the sizeable whole-mount specimen was the refractive index mismatch that occurred at phase boundaries in the observation setup. Because the point spread function tells that the magnitude of image distortions is exponentially proportional to the length of the light path, the refraction at the phase boundaries was carefully eliminated from the whole light path, especially during the observation of the volumetric 3D sample.

Mounting a biological specimen in the TDE-filled chamber, as described above, eventually provides refractive index mismatch-free preparation of embryonic tissues for 3D observation. As our study has demonstrated, it is possible to obtain a clear and bright image from never before reached deep inside of specimens with a conventional confocal microscope.

### 5. Whole-mount in situ hybridization

The embryos were fixed in 4%PA in 0.1M PB overnight and processed for whole-mount in situ hybridization, as described by Homma et al. (2007).

## ACKNOWLEDGEMENT

The corresponding author would like to dedicate this paper to Dr. Ronald W. Oppenheim. I am so grateful that he took me as a student when I started my academic carrier, and I will be forever proud to have him, my mentor.

